# Comparative genomics reveals evolutionary drivers of sessile life and left-right shell asymmetry in bivalves

**DOI:** 10.1101/2021.03.18.435778

**Authors:** Yang Zhang, Fan Mao, Shu Xiao, Haiyan Yu, Zhiming Xiang, Fei Xu, Jun Li, Lili Wang, Yuanyan Xiong, Mengqiu Chen, Yongbo Bao, Yuewen Deng, Quan Huo, Lvping Zhang, Wenguang Liu, Xuming Li, Haitao Ma, Yuehuan Zhang, Xiyu Mu, Min Liu, Hongkun Zheng, Nai-Kei Wong, Ziniu Yu

## Abstract

Bivalves are species-rich mollusks with prominent protective roles in coastal ecosystems. Across these ancient lineages, colony-founding larvae anchor themselves either by byssus production or by cemented attachment. The latter mode of sessile life is strongly molded by left-right shell asymmetry during larval development of *Ostreoida* oysters such as *Crassostrea hongkongensis*. Here, we sequenced the genome of *C. hongkongensis* in high resolution and compared it to reference bivalve genomes to unveil genomic determinants driving cemented attachment and shell asymmetry. Importantly, loss of the homeobox gene *antennapedia* (*Antp*) and broad expansion of lineage-specific extracellular gene families are implicated in a shift from byssal to cemented attachment in bivalves. Evidence from comparative transcriptomics shows that the left-right asymmetrical *C. hongkongensis* plausibly diverged from the symmetrical *Pinctada fucata* in expression profiles marked by elevated activities of orthologous transcription factors and lineage-specific shell-related gene families including *tyrosinases*, which may cooperatively govern asymmetrical shell formation in *Ostreoida* oysters.

## Introduction

Bivalves belong to the ancient lineages of *Mollusca* comprising nearly 9,600 species that thrive in aquatic environments, with notable economic and ecological importance [1]. As bilaterian organisms, they rely nutritionally on filtering phytoplankton, and primarily follow a life cycle that transitions from free-swimming larvae to attached juveniles, culminating in sessile life [2, 3]. Among filter-feeding bivalves, oysters of the superfamily *Ostreoidea* serve as crucial guardians of marine ecosystems by forming oyster reefs that clean up water and sustain biodiversity [4,5]. Due to climate change and coastal degradation, however, bivalves face profound challenges from warming waters and ocean acidification, which destabilize habitats, raise infection risks and dampen the bivalve capacity of acquiring carbonate for shell formation [6–8].

To cope with diverse ecosystems, a variety of sessile strategies has emerged in bivalves during evolution, among which two modes of sessile life prevail. Characteristically, majority of the bivalves, including *Mytilidae* (mussel), *Pectinidae* (scallop), and *Pteriidae* (pearl oyster) secret adhesive byssal threads to stabilize themselves against marine turbulences [9–13]. In contrast, *Ostreoida* oysters have evolved a highly sophisticated machinery of cemented attachment through producing organic-inorganic hybrid adhesive substances in place of byssus, which allows them to permanently fuse the left shell with rock surfaces or shells of other individuals in intertidal zones [14]. Compared with byssus, cemented attachment exhibits superiority in physical adhesion and mechanical tension, enabling oysters to efficiently create and thrive in large reef communities [2]. Developmentally, as a salient feature of their exoskeleton, shell formation processes in bivalves are strongly molded by their preferences for sessile life [15]. Quite distinctively, byssally attached bivalve species tend to possess a bilaterally symmetrical shell, whereas cement-attached oysters present a high degree of phenotypic variability and morphological asymmetry characteristic of their radically distinct left-right (L/R) shells [15]. Nevertheless, the molecular mechanisms driving these extraordinary innovations in bivalve evolution remain enigmatic, particularly in genomic contexts.

The Hong Kong oyster (*Crassostrea hongkongensis,* first described as *Crassostrea rivularis* by Gould, 1861) is an economically valuable aquacultural species endemic to the South China coastline [16]. As an ideal model for studying shell asymmetry, *C. hongkongensis* larvae follows a typical developmental cycle of cemented attachment and asymmetrical differentiation of the L/R shells. In order to elucidate the genetic basis underpinning the evolution of bivalve sessile life and asymmetry of shell formation, we sequenced and analyzed the complete genome of *C. hongkongensis* and performed comparative genomic analysis along with several other bivalve species, including two congeneric *Ostreoida* oysters, *Crassostrea gigas*, and *Crassostrea virginica* [12,17–21]. In addition, we monitored transcriptomic changes of *C. hongkongensis* embryos during the critical window of larval attachment, and compared any asymmetry-related gene expression patterns in the L/R mantles of adult *C. hongkongensis* and byssus-producing pearl oyster (*Pinctada fucata*). Our comparative genomic data and associated functional assays reveal extensive molecular adaptations across the oyster genome that support the evolutionary switch from byssal to cemented attachment and divergence from symmetrical shell in *Ostreoida* oysters.

## Results

### Genome sequencing, annotation and Hi-C, phylogenomics and evolutionary rate

Efforts on genome sequencing and assembly are inherently challenging for many marine invertebrates such as mollusks, annelids, and platyhelminths due to their remarkable genetic heterozygosity (or polymorphisms) [17,18,21,22]. Based on k-mer analysis, the genome size of a single wild-stock Hong Kong oyster (*C. hongkongensis*) individual was estimated to be 695 Mb with 1.2% of heterozygosity (**Figure S1**), which is broadly comparable to that of the Pacific oyster (1.3%) [17]. To circumvent limitations of short-read next-generation sequencing in assembling highly polymorphic genomes, PacBio sequencing in combination of Illumina sequencing was instead opted as the dominant mode of genome sequencing in our study. We first generated 23.25 Gb of raw PacBio reads and 147.25 Gb of Illumina reads, being equivalent to 31.9-fold and 201.8-fold genome coverage, respectively (**Table S1 and S2**). Following stepwise optimization of assembly algorithms, these reads were assembled into a 729.6 Mb genome with a contig N50 of 314.1 kb and a scaffold N50 size of 500.4 kb, with the longest contig spanning 2.37 Mb (**Table S3**). The contig N50 of the oyster genome is at least one order of magnitude more expansive than those of published bivalve genomes (**Table S4**), demonstrating the superiority of long-read sequencing technologies in coping with high polymorphism in genome assembly of marine invertebrates. However, the assembled genome size turned out to be slightly larger than that estimated by k-mer analysis. Such discrepancy may reflect sequence preferences of Illumina reads. The high integrity and quality of the assembly were evidenced by a productive mapping of 97.57% of sequencing reads and a low single-nucleotide error rate (**Table S5 and S6**). Moreover, Benchmarking Universal Single-Copy Orthologs (BUSCOS) analysis confirmed a high degree of completeness (92.84%) for the assembled genome (**Table S7**), which is comparable in genome completeness to other published bivalves (**Table S4**).

In order to assemble the oyster genome to chromosomal level, we generated ~44.4 million valid Hi-C interaction pairs with over 50-fold coverage (**Table S8**). Meanwhile, 690.39 Mb of genome sequence were anchored into 10 of pseudo-chromosomes with Hi-C data by using LACHESIS, covering 94.66 % of the assembled genome (**Figure 1A**, **Figure S2** & **Table S9**). Among them, 648.56 Mb of genome sequence were reoriented and anchored into chromosomes, constituting 93.94% of the total anchored sequences (**Table S9**). Moreover, high consistency between Hi-C based pseudo-chromosomes with the genetic map of one congeneric species, *C. gigas*, was confirmed (p = 0.978-0.996, **Figure S3**), implicating high reliability in chromosomal genome assembly. Overall, by leveraging PacBio and Hi-C enhanced Illumina sequencing, a very high quality and chromosome-anchored complete genome was obtained, thus providing a robust framework for subsequent exploration of oyster biology and evolution of bivalves.

**Figure 1.**
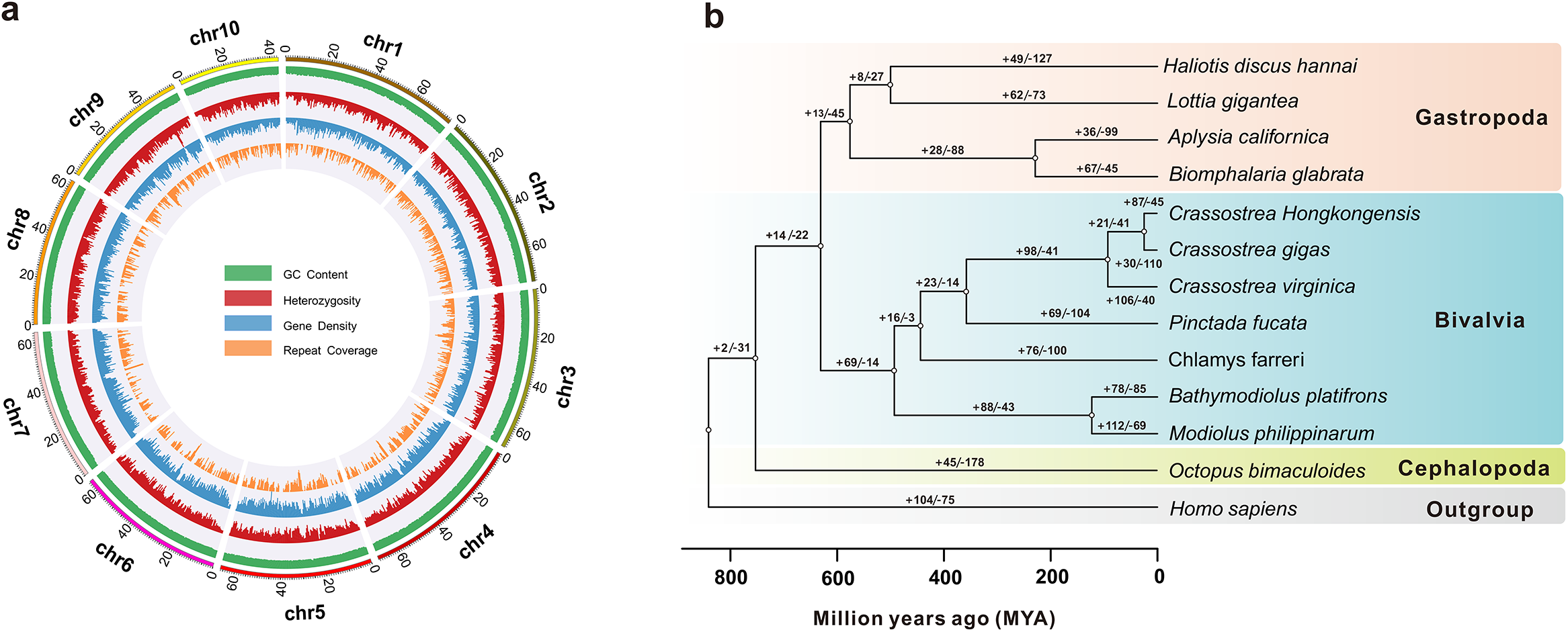
The genome landscape and phylogenetic analysis of the oyster *Crassostrea hongkongensis*. **A.** Circos plot highlights genome characteristics across 10 chromosomes in a megabase (Mb) scale. The GC content, global heterozygosity, gene density and repeat coverage are presented from outer to inner circles in turn with non-overlapping 1 Mb sliding windows. **B.** Analysis on gene family expansion/contraction and divergence time across 12 representative mollusks species. A total of 87 gene families are expanded in the Hong Kong oyster, *C. hongkongensis*. The human genome was set as an outgroup. Three *Ostreoida* oyster species (*Crassostrea hongkongensis*, *Crassostrea gigas*, *and Crassostrea virginica*) are clustered together. Gene family expansion/contraction is indicated by a plus or minus sign.

For gene annotation, we predicted 30,021 protein-coding genes in the genome by integrating results from *ab initio* prediction, homology-based searches with reference genomes and RNA-seq (**Table S10**), with an estimated BUSCO completeness of 91.09% (**Table S11**). Of these, more than 97.97% of the predicted genes (28,329 genes) were annotated in the public databases (**Table S12**). The gene number here resembles that in a close relative species, *C. gigas* (28,027) [17]. In addition, transposon elements (TE) constitute 46.2% of the *C. hongkongensis* genome, among which the prevailing TE is class II Helitron (12.4%, 90.4 Mb) (**Table S13**). Phylogenetic analysis showed that three *Ostreoida* oyster species (*C. hongkongensis*, *C. gigas*, *C. virginica*) clustered together (**Figure 1B**), and that *Ostreoida* oyster speciation took root around 92.1 million years ago (Mya), in agreement with evidence from mitochondrial genomes [23]. Within bivalves, *Ostreoida* oysters are closest to the *Pteriidae* oyster *Pinctada fucata*, and their point of divergence was estimated to be 357.5 Mya (Figure 1B). These results corroborates the hypothesis that a common ancestor of primitive *Ostreoida* and *Pteriidae* oysters existed prior to the Permian-Triassic extinction event, whereas speciation of modern *Ostreoida* oysters began at the end of Cretaceous-Paleogene extinction event [24–27]. Consistently, comparative genomic synteny shows high genomic collinearity between three *Ostreoida* oyster genomes except for large intra-chromosomal inversions, but substantial inter-chromosomal translocations and rearrangements occur between chromosomes of *Ostreoida* oysters and *Pinctada fucata* (**Figure S4**), which is in agreement with their phylogenetic relationship and duration of divergence.

### *Homeobox* gene cluster

Radical changes toward a sessile life require evolutionary innovations in anatomical organization. In contrast to byssus-producing bivalves [12,28], *Ostreoida* oysters do not possess a byssal gland or secret byssus during lifetime [29,30], though a vestigial foot transiently appears at the veliger stage and degenerates following attachment and metamorphosis (**Figure 2B**). Developmentally, the *homeobox (Hox)* genes are known for their crucial roles in regulating body-plan development and organogenetic transitions in metazoans [31–34]. In view of this, we compared the clustering of *Hox* genes in byssus-producing and byssus-null bivalve species. A salient feature in byssal bivalves including *Pinctada fucata*, *Mizuhopecten yessoensis*, *Chlamys farreri*, *Mytilus galloprovincialis*, *Bathymodiolus platifrons*, and *Modiolus philippinarum* is an intact *Hox* and *para-Hox* gene cluster (**Figure 2A** and **Figure S5**). In contrast, a disputed *Hox* gene cluster reportedly exits in *C. gigas* oyster genome [17], whereas a coherent *Hox* gene cluster is configured linearly in one single-locus in both *C. hongkongesis* and *C. virginica*, probably in part due to fragmented genome assembly in *C. gigas*. Intriguingly, one of the key *Hox* members *antennapedia* (*Antp*) is lost in all three *Ostreoida* oysters (Figure 2A), thereby implicating *Antp* gene as an essential driver of byssus formation. Sequence alignment reveals that *Antp* gene possesses a conserved homeobox domain in bivalves (**Figure S6**).

**Figure 2.**
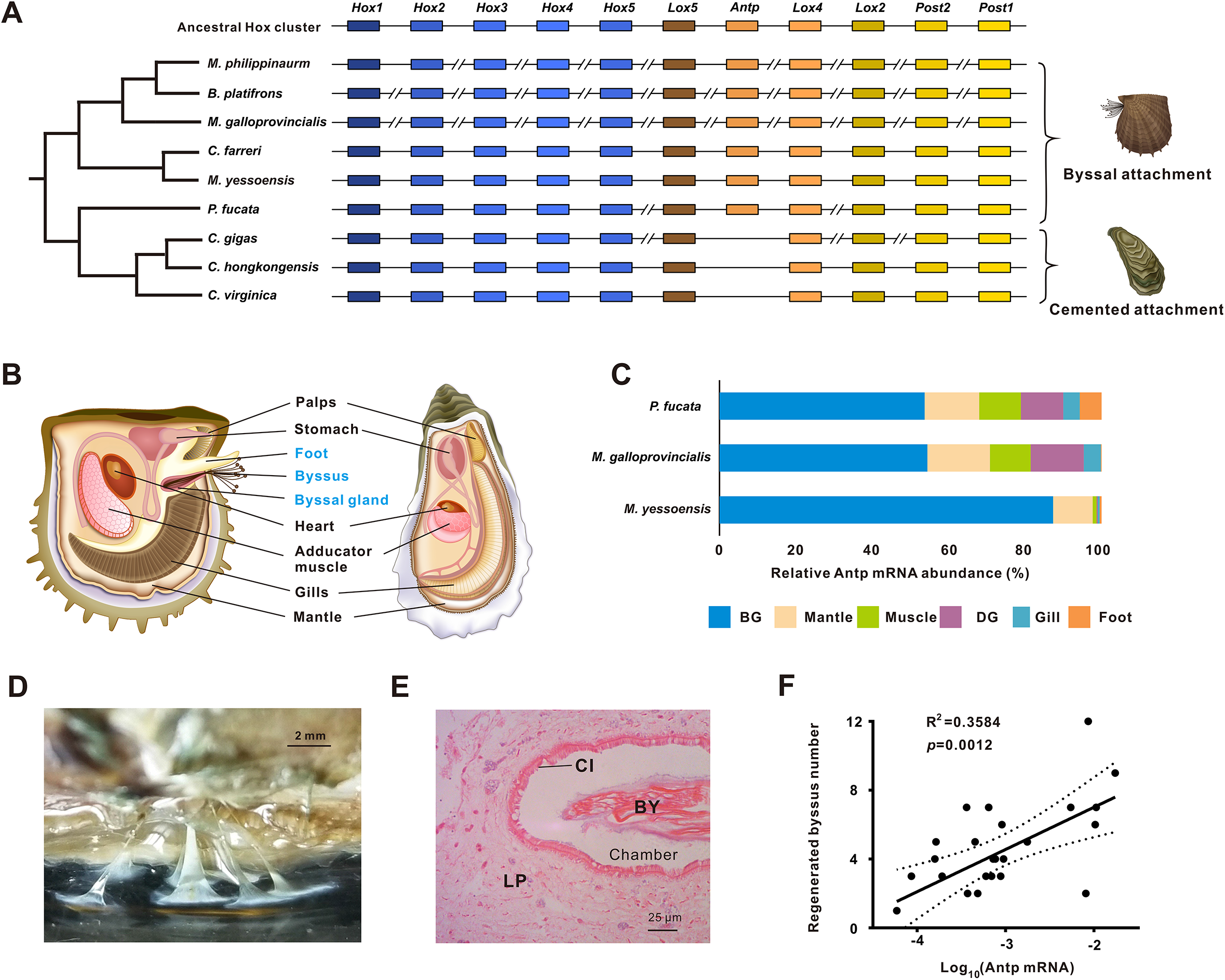
Loss of the homeobox gene *antennapedia (Antp)* is implicated in an adaptive shift from byssal attachment to cemented attachment. **A.** Comparison of *Homobox* (*Hox*) cluster organization in bivalves with two distinct attachment styles, byssal attachment and cemented attachment. Unlike the disputed *Hox* gene cluster in *C. gigas* oyster genome, *Hox* gene cluster configures linearly in both *C. hongkongesis* and *C. virginica*. Essentially, *Antp* is lost in all three *Ostreoida* oysters. **B.** Overview of key body-plan organization in *Pinctada fucata* and *C. hongkongensis*. *P. fucata* possesses a byssal gland and byssus, whereas adult individuals of *Ostreoida* oyster have lost their byssus gland and byssus. **C.** Tissue distribution of *Antp* othologues in three byssally attached bivalves, *P. fucata*, *Mytilus galloprocincialis*, and *Mizuhopecten yessoensis*. BG, byssal gland; DG, stomach. *Antp* mRNA abundance is displayed in percentage, and its expression in BG accounted for more than 50%. **D.** Morphology of newly regenerated byssus 48 h after excision of original byssus. Scale bar: 2 mm. **E.** Anatomic analysis of byssal gland with cross section. Vertical cross-section of the byssal gland displaying ciliated walls (Cl), lamina propria (LP), and byssal remnants (BY) within a chamber. Scale bar: 50 μm. **F.** Correlation between abundance of *Antp* mRNA and regenerated byssus numbers in *P. fucata*. *Antp* mRNA level in the byssal gland was determined by real-time qPCR, while newly regenerated byssus threads was counted 48 h after excision of original byssus. Pearson’s correlation coefficients and *p*-values were calculated with two tailed tests with 95% confidence.

As evidenced in expression profiles of three representative byssus-producing bivalve species, *Antp* and its orthologues are predominantly expressed in the byssal gland (**Figure 2C**). Due to unavailability of molecular tools like CRISPR/Cas9 or TALEN for manipulating bivalve genomes, genetic ablation of the *Antp* gene is not yet feasible in pearl oyster for phenotypic appraisal of its function. However, histological evidence suggests that the byssal gland is one of the appendage organs capable of secreting thin extended byssal threads in their mature form as observable bysuss outside the organism (**Figure 2D and Figure 2E**). Based on the fact that regenerative ability varies among individuals, we assessed *Antp* function in this phenotypic trait. Remarkably, mRNA expression levels of *Antp* are highly correlated with the number of regenerative byssus in the pearl oyster (*n* = 24, *R*^2^ = 0.36, *p* = 0.0012; **Figure 2F**). Taken together, our evidence strongly implicates *Antp* as a transcriptional regulator central to byssal secretion in *P. fucata*. Further, the loss of the *Antp* gene seems to be associated with a physical loss of byssal gland in oysters. In an evolutionary perspective, *Antp* seems to play a critical role in appendage diversification in arthropods, which has previously been evidenced by its involvement in leg formation in the crustacean *Daphnia* [35], and repression of abdominal limb in the spider *Achaearanea tepidariorum* [36]. In addition, ectopic expression of *Hox* transcription factor *Antp* reportedly induced expression of the silk protein sericin-1 as a biopolymer in the silkworm *Bombyx mori* [37,38]. Collectively, these findings support a conserved function of *Antp* in secretory appendage in two distinct lineages, mollusks and arthropods.

### Gene expansion and oyster attachment

In place of byssal attachment, *Ostreoida* oysters adopt an ingeniously cost-effective way of sessile life, namely, cemented attachment [29,30]. Such adhesive mechanism is characterized by extraordinary mechanical strength and superior flexibility needed to resist powerful tidal scour and absorb surge energy [39]. Cemented attachment allows oysters to efficiently anchor and thrive in marine environments, and ultimately supports the genesis and health of oyster reefs. Nevertheless, the molecular mechanisms underlying oyster adhesive production have remained enigmatic. Taking into account that commented attachment is an innovation unique to *Ostreoida* oysters, we first ventured to investigate which gene families are expanded as a common event in three *Ostreoida* species. Our results show that in *C. gigas*, *C. hongkongensis*, and *C. virginica*, there are 58, 172, and 321 expanded species-specific gene families, respectively, which can be further reduced to 32 expanded core gene families in *Ostreoida* oyster genomes (**Figure 3A** and **Figure S7 & Table S14**).

**Figure 3.**
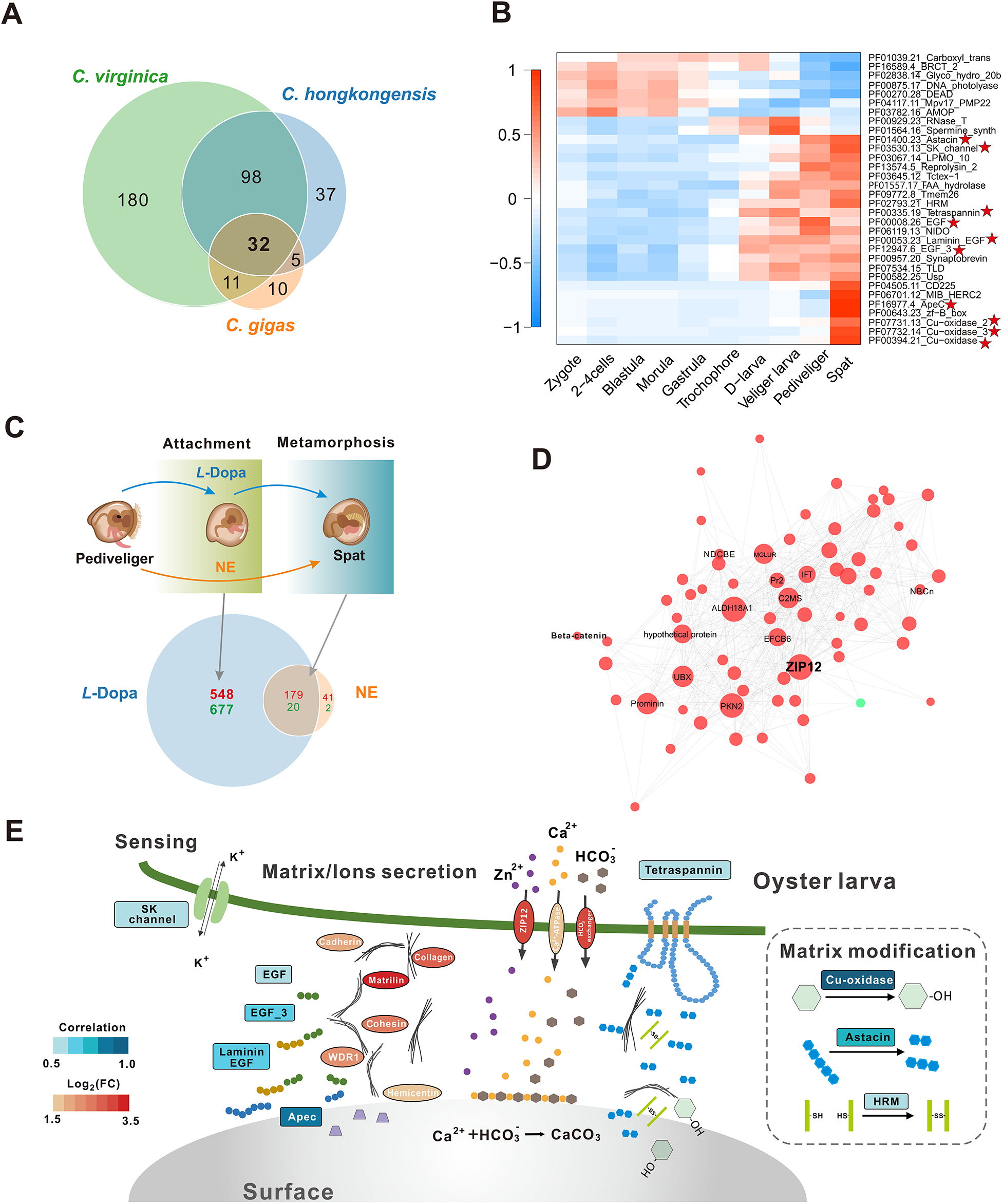
Molecular basis of attachment initiation in *Ostreoida* oysters. **A.** Veen plot shows the common gene family expansion in three *Ostreoida* oyster species, *C. hongkongensis, C. virginica*, and *C. gigas*, among which 32 core gene family expansions were identified. **B.** Heatmap illustrates the correlation between expression levels of 32 core gene family and developmental stages of *C. hongkongensis* larvae. The high correlation of transcriptional activated gene family with attachment is presented by red. **C.** Pharmacological responses of oyster veliger larvae during attachment initiation and metamorphosis. *L*-3,4-dihydroxyphenylalanine (*L*-DOPA) stimulated larval attachment and metamorphosis, while noepinephrine (NE) only induced metamorphosis without attachment. Veen plot shows that *L*-DOPA/NE induced specific genes, among which *L*-DOPA specifically induced genes may participate in attachment initiation. **D.** Construction of coordinated gene networks based on the zinc transporter *ZIP12*, which is a hub forming the highest degrees of gene connections in Weighted correlation network analysis (WGCNA) analysis. Red and green dots indicate up-regulated genes and down regulated genes. **E.** Schematic diagram conceptualizing the molecular basis for initiation of larval attachment in oysters. Square box indicates oyster-specific expanded gene families involved in larval attachment (*p* <0.001). Filled color (blue) was scaled with correlation values at the spat stage. Ellipse box indicates *L*-DOPA specifically induced genes after *L*-DOPA treatment, which are filled in red scaled with values in log_2_ (FC). FC, fold change.

To elucidate how expansion of these core gene families facilitates cemented attachment, we determined the correlations between their expression levels and specific developmental stages (**Figure 3B**). Developmentally, attachment is an intricate secretion process involving a broad spectrum of chemical reactions and proteins, notably extracellular enzymes or matrices [40]. It is thus unsurprising to identify a *small conductance calcium-activated potassium channel* (*SK channel*) gene family and 9 extracellular gene families at work in this process, which show high correlations in the pediveliger and spat stages corresponding to larval initiation of attachment. *SK channels* are widely expressed calcium-activated potassium channels in neurons [41,42], with crucial roles in regulating dendritic excitability, synaptic transmission, and synaptic plasticity [43,44]. Interestingly, increased expression of expanded *SK channels* may aid free-swimming larvae in sensing external environments in search for an appropriate attachment site. On the other hand, the function of extracellular gene families is strictly related to key processes of shell attachment, including matrix secretion (*Epidermal growth factor (EGF)*, *EGF3*, *lamin EGF*, *Apec*), processing of matrix modification (*Cu-oxidase*, *Cu-oxidase2*, *Cu-oxidase3*, and *astacin*), among others (Figure 3B). Indeed, many adhesive proteins contain specific protein-binding domains [45], such as EGF-like domains in the slug mucus proteins (e.g. Sm40 and Sm85) [46] and sea star footprint proteins (e.g. Sf1) [47], raising the possibility that *EGF* family expansion in *C. hongkongensis* is functionally linked to cemented attachment. Additionally, physico-chemical properties of many adhesive proteins arise in part from post-translational modifications, which ultimately support their adhesive functions [45]. Protein oxidation in marine bio-adhesives indeed contributes to enhanced crosslinking between shell disks and substrates during attachment [48,49]. A notable gene expansion in the copper oxidase family is likely to contribute to stabilization of extracellular matrixes in the form of crosslinking between the oyster shell and external substrates. *Copper-based enzyme lysyl oxidase* is known to be essential for cross-linking and strengthening fibers in animal connective tissues via collagen oxidation [50]. Concomitantly, copper ion, as part of oxidative enzymes, is a mandatory cofactor for oxidase activity, which creates cross-linking sites from common amino acids, to enhance the cemented attachment [51–53]. In further transcriptomic analysis, we found evidence that 9 extracellular gene families were starkly upregulated during the larvae-spat transformation of embryo development stages (**Figure S8**), corroborating their functional importance in attachment formation.

### *L*-DOPA induced attachment

During larvae-spat transformation, embryonic oysters execute an intrinsic program of developmental changes, in which cemented attachment is tightly coupled to metamorphosis [54]. In this context, we set out to distinguish molecular determinants of cemented attachment from that of metamorphosis at the veliger stage by means of two pharmacologic agents: *L*-3,4-dihydroxyphenylalanine (*L*-DOPA) and norepinephrine (NE) at the veliger stage. The former simultaneously promoted normal attachment and metamorphosis, whereas the latter induced metamorphosis only but not attachment (**Figure 3C** and **Figure S9**) [54]. Based on this, gene expression induced by *L*-DOPA rather than NE was hypothesized to be a driver for the initiation of attachment in *C. gigas*. We accordingly scrutinized 24 transcriptomes following pharmacological challenges at two time points within the temporal span of oyster attachment. Our results show that the expression of 1225 genes was specifically altered by treatment of *L*-DOPA rather than NE (Figure 3C), confirming the former’s essential roles as an attachment signal.

Remarkably, several neurotransmitter receptors (including *metabotropic glutamate receptor* and *neuropeptide Y receptor*) were starkly increased, consistent with the assumption that neuromuscular coordination is mandatory for guiding embryos to settle in suitable niches and initiate attachment (**Figure S10**) [55,56]. Moreover, genes of metal ion channels or binding proteins were significantly enriched, with notable examples like organic cation transporter protein, *transient receptor potential cation channel* (*ZIP12*) and *voltage-dependent calcium channel* (*Ca^2+^-ATPase*), which is intuitively consistent with the well-documented stimulatory roles of selective cations in oyster larval settling [57]. To highlight, potassium voltage-gated channel activity was proven to be vital for oyster larval attachment, since its inhibitor tetraethyl ammonium can effectively block this developmental process [58]. Typically, attachment initiates in oyster larvae with the aid of fibrous adhesive proteins and other bioorganic substances including mucopolysaccharides and phospholipids [2,59]. As a consequence, extensive extracellular matrix and adhesion proteins including collagen, cadherin, fibrocystin, and hemicentin would increase in response to *L*-DOPA simulation, presumably paving the way for larval attachment [60].

To search out the crucial molecular determinants governing this process, we performed WGCNA to construct a potential connected gene network functionally associated with *L*-DOPA induced attachment, wherein 15 of modules were subsequently identified (**Figure S11**). Among them, the MEpink module is the most correlated with *L*-DOPA induced attachment (*p* < 0.01) and contains 139 of genes (topological overlap > 0.3). Intriguingly, within this module, a hub forming the most connections in the network was found to be zinc transporter *ZIP12* (**Figure 3D**), which is a pivotal regulator of zinc flux. As a co-factor essential to a wide spectrum of proteins such as matrix metalloproteinases, zinc plays vital regulatory roles in enzymatic catalysis and macromolecular stability [61]. High abundance of zinc is also a salient feature in aragonite- or calcite-rich shells in certain mollusks [62]. Meanwhile, among the gene families that specifically expanded in *Ostreoida* oysters, *astacin* is a cell-secreted or plasma membrane-associated protease that possesses zinc binding activity and takes part in proteolytic processing of extracellular proteins [63]. Its expression was markedly elevated both during larvae-spat transformation or larval response to *L*-DOPA treatment (**Figure S8g & S10d**). Predictably, chelation of zinc potently retarded oyster larval attachment (**Figure S12**), providing additional hints that initial creation of matrix structures requires zinc and associated protein activities for cement attachment. Accordingly, based on genomic results on extracellular gene family expansion and transcriptomic profiles for the attachment stage, we conceived a conceptual model to delineate the mechanistic determinants and processes at work in the cement attachment strategy of oyster larvae (**Figure 3E**). We postulate that attachment formation apparently results from an intricate coordination of at least three types of fundamental activities, namely: larval sensing of habitable surfaces, matrix/ion secretion, and matrix modification to mobilize adhesive processes.

### Asymmetry in left-right shell formation

Symmetry is an elegant guiding principle for the implementation of body plans [64]. Across the class *Bivalvia*, majority of bivalves display a perfect or near-perfect conformity to bilaterally symmetrical shells [15,65]. In contrast, *Ostreoida* oysters may appear unorthodox in adopting morphological asymmetry in their shell formation due to functional differentiation of the left-right (L/R) shells (**Figure S13**). The left shell is visibly much thicker and more convex than its right counterpart, which is apt for attaching to rocky surfaces or neighboring oysters within a reef community. On the other hand, the right shell is capable of physical displacement and hermetic lockdown to regulate water intake and ward off predation (**Figure 4A**). Moreover, structural variance in shell asymmetry is also amply reflected by a greater proportion of prismatic layer in the right shell (**Figure S14**), which is responsible for controlling initiation of calcite crystal formation and growth [66,67]. Although asymmetry of body forms has been traditionally stereotyped as defects that may jeopardize survival of an organism [68], the example of *Ostreoida* oysters clearly defies this rule. We reason that such an intriguing differentiation of asymmetrical shells could confer unexpected benefits such as improved population fitness in an otherwise intrinsically harsh coastal environment. With the advent of the left shell and its versatile attachment machinery, oysters can easily economize resources or secure their foothold on rocks or peers’ shells within an oyster reef via cemented attachment [69]. This strategy permits oysters to lower their thresholds for founding and expanding productive colonies in demanding physical habitats, literally through stacking of individuals at high densities, without sacrificing resistance to environmental challenges such as tidal turbulences.

**Figure 4.**
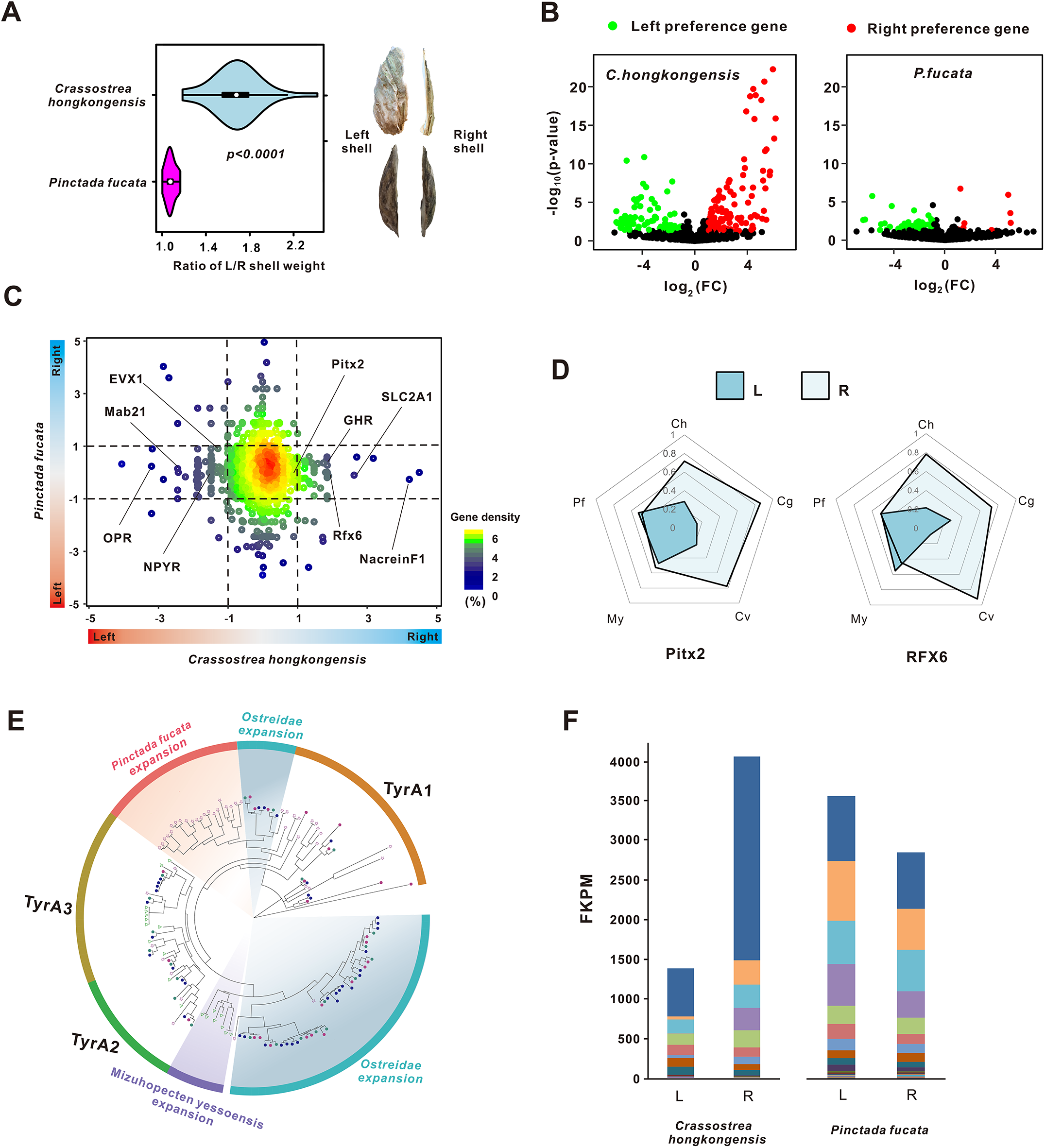
Left-right asymmetry of shell formation in *Ostreoida* oysters. **A.** Comparison of the ratio of left/right shell weight and morphology between the *C. hongkongensis* and *P. fucate*. **B.** Volcano plot shows the left- and right-mantle differentially expressed genes, which are filtered by |log_2_(FC)| ≥1 with *p*-value <0.05. **C.** Expression profile of 1:1 orthologues in L/R mantle of *C. hongkongensis* and *P. fucate*. A total of 10,491 orthologues were paired and only a few asymmetrical orthologues were specifically expressed in the Hong Kong oyster. The *x*- and *y*-axes indicate logFC of expression ratio in R/L mantle of *C. hongkongensis* and *P. fucate*, respectively. **D.** Expression patterns of two pivotal transcription factors in left and right mantles across five bivalves. Ch, *C. hongkongensis*; Cg, *C. gigas*; Cv, *C. virginica*; My, *M. yessoensis*; Pf, *P. fucate*. **E.** Dendrogram of known *tyrosinases* from five mollusks was constructed by maximum likelihood (ML) method. Bivalve and molluscan TyrA orthologous groups are indicated by curvatures and annotated as A1-A3. Specific *tyrosinase* orthologous groups are marked with color background and annotated with a species’ name. Species are represented with different shapes: triangle, *Mizuhopecten yessoensis*; circle, *Ostreidae*; pentagon, *Pinctada fucata*. **F.** Expression patterns of *tyrosinase* families in left and right mantles of two bivalve species, as determined by FPKM. Total FPKM of different types of orthologous genes is displayed in cumulative histograms. Different members of *tyrosinase* are presented with different colors.

To further elucidate the molecular basis of left-right asymmetry, comparative transcriptomics was carried out for quantify the gene expression profiles in L/R mantles of *C. hongkongensis* and pearl oyster, which are the key organ controlling shell formation [70,71]. As expected, 188 asymmetry-related differentially expressed genes (DEGs) of the L/R mantles were identified in *C. hongkongensis*, whereas only 53 asymmetry-related DEGs were found in the pearl oyster (**Figure 4B**), which reflects a radical genetic divergence underpinning shell asymmetry. Next, to test the hypothesis that lineage-specific divergence of orthologues contribute to symmetry breakage, 10,050 of the orthologues were paired between the two species (**Table S15**). Our results indicate that a few but crucial asymmetry-related orthologues are specifically expressed *C. hongkongensis* (**Figure 4C**), including homeobox gene *paired-like homeodomain transcription factor* (*Pitx2*) and *homeobox B4a* (*Hox-B4a*), and *regulatory factor X6* (*RFX6*). Notably, *Pitx2* is a central regulator orchestrating the Nodal cascade, which is responsible not only for directing L/R axis formation in mammals [72], but also shell coiling and L/R asymmetry in some mollusks such as the snail [73]. Another gene of interest is the *RFX6*, recognized for its fundamental importance in guiding pancreatic islet development and insulin production in mammals [74]. While insulin-related peptide gene is known for being a critical driver of oyster growth [75], this new evidence alludes to novel roles of Rfx6-insluin signaling in maintaining shell asymmetry in oysters. As predicted, asymmetry-related expression of *Pitx2* and *RFX6* in L/R mantles was confirmed by real-time qPCR in three *Ostreoida* lineages with asymmetrical shells, whereas such gene expression patterns were absent in three symmetrical bivalves, pearl oyster, scallop and mussel (**Figure 4D**).

However, it should be noted that majority of asymmetry-related genes in *C. hongkongensis* are not orthologous to the pearl oyster. For example, *tyrosinases* are one of the key gene families involved in steering shell formation and pigmentation by means of oxidation and cross-linking of o-diphenols [76,77]. Phylogenetic analysis reveals that more than a half of *tyrosinase* genes (55%) clustered in several lineage-restricted clades, suggesting rapid and independent expansion of this gene family in bivalves (**Figure 4E**). Remarkably, several high-abundance members of the *tyrosinase* family seem to be strongly associated with L/R asymmetry and were expressed preferentially in the right mantles of *C. hongkongensis*, whereas no obvious variance was noted between L/R mantles in the pearl oyster (**Figure 4F**). Therefore, it seems logical to infer that rapid expansion and divergent expression of *tyrosinase* family contribute importantly to the emergence and neofuncationalization of asymmetrical shell formation in *Ostreoida* lineages. Lastly, in determining when precisely expression of these asymmetry-related genes kick off in oyster embryogenesis, we found that 71.1% of these genes start expression at the spat stage (**Figure S15**), implying that a complete asymmetrical pattern becomes established in the juveniles only after metamorphosis.

## Conclusion

*Ostreoida* oysters have evolved remarkable innovations for streamlining their bodyplans, which are enabled by novel cemented attachment and an allied gene machinery diverging from L/R symmetry. These evolutionary breakthroughs poise oysters as highly successfully reef builders and ecological guardians integral to marine ecosystems spanning the globe. To reveal the genomic changes driving these evolutionary innovations, we sequenced the complete genome of *C. hongkongensis*, obtained active transcriptomes developmentally critical to the attachment window, and made comparisons with other bivalve genomes. The homeobox gene *Antp* of the Hox cluster, found to be lost in *Ostreoida* oysters, is evidently a pivotal regulator of byssal secretion and expression of byssal proteins in *P. fucata*, and potentially a critical gene governing the radical switch from byssal to cemented attachment. Furthermore, extensive extracellular gene families were expanded in the *Ostreoida* lineages specifically, presumably contributing to the operationalization of cemented attachment. Ion-binding genes were significantly enriched in *L*-DOPA induced attachment in oyster, with zinc-binding genes being a prominent network that coordinates extracellular matrix modification and initiates adhesion. Moreover, *Ostreoida* divergence from shell symmetry is probably under the joint control of a suite of transcriptionally identified asymmetry-related DEGs of the L/R mantles, notably the transcription factors *Pitx2* and *RFX6*, as well as expanded lineage-specific family of *tyrosinases*. Thus, on the basis of genomic determinants and coordinated gene networks as revealed in this study, we have advanced a detailed picture of how shell asymmetry is switched on and driven in bivalves such as *Ostreoida* oysters. In order to provide insights into bivalve biology and disease in contexts of climate change or biological conservation, further investigation on the attachment-governing genes may be warranted.

## Materials and methods

### Illumina sequencing

Genomic DNA was extracted by using DNeasy Blood & Tissue Kit (Cat. no. 69582, Qiagen, Germany) from a two-year old single individual of *C. hongkongensis*. Two types of pair-end libraries (220 bp and 500 bp) and six types of long-insert mate-pair libraries (3 kb, 4 kb, 5 kb, 8 kb, 10 kb, and 15 kb) were constructed by using Illumina’s paired-end and mate-end kits, according to the manufacturer’s instructions. Libraries were sequenced on an Illumina Hiseq 2500 platform. For raw reads, sequencing adaptors were removed. Contaminated reads (such as chloroplast, mitochondrial, bacterial, and viral sequences, etc.) were screened by alignment in accordance with an NCBI-NR database by using BWA v0.7.13 [78] with default parameters. FastUniq v1.1 [79] was used to remove duplicated read pairs. Low-quality reads were filtered out, according to the following criteria: 1) reads with ≥10% unidentified nucleotides (N); 2) reads with >10 nucleotides aligned to an adapter, allowing ≤10% mismatches; 3) reads with >50% bases with Phred quality <5.

### PacBio sequencing

Genomic DNA was sheared by a g-TUBE device (Cat. no. 520079, Covaris, MA) with 20 kb settings. Sheared DNA was then purified and concentrated with AMPure XP beads (Cat. no. 10136224, Beckman Coulter, CA) and further used for single-molecule real-time (SMRT) bell preparation according to the manufacturer’s protocol (Pacific Biosciences, CA), and 20 kb template preparation by using BluePippin size selection (Sage Science). Size selected and isolated SMRT bell fractions were purified with AMPure XP beads. Finally, these purified SMRT bells were used for primer and polymerase (P6) binding, according to manufacturer’s binding calculator (Pacific Biosciences). Single-molecule sequencing was performed on a PacBio RS-II platform with C4 chemistry. Only PacBio subreads no shorter than 500 bp were included for performing oyster genome assembly.

### Genome size estimation

About 34 Gb (52×) corrected Illumina reads from the 180 bp and 500 bp were selected to perform genome size estimation. The oyster genome size was estimated based on the formula: Genome size = Kmer number/Peak depth.

### *De novo* genome assembly of Illumina data

Clean Illumina reads were assembled *de novo* into longer contigs by using ALLPATH-LG [80] with default parameters. Adjacent contigs were linked to scaffolds by leveraging mate-pair information with SSPACE v2.3 [81], while gaps were filled by using GapCloser v1.12 [81] implemented in a SOAPdenovo2 package [82].

### *De novo* genome assembly of PacBio data

#### Canu+LoRDEC+WTDBG

We used an error correction module of Canu v1.5 [83] to select longer subreads with the settings ‘genomeSize = 3,500,000,000’ and ‘corOutCoverage = 80’, detect raw subreads overlapping through a highly sensitive overlapper MHAP v2.12 (‘corMhapSensitivity = low/normal/high’), and complete an error correction through a falcon_sense method (‘correctedErrorRate = 0.025’). Subsequently, output subreads of Canu were further corrected by LoRDEC v0.6 [84] with the parameters ‘-k 19 -s 3’. Based on these two rounds of error-corrected subreads, we generated a draft assembly by using WTDBG 1.1.006 (https://github.com/ruanjue/wtdbg) with the command ‘wtdbg -i pbreads.fasta -t 64 -H -k 21 -S 1.02 -e 3 -o wtdbg’.

### Hybrid genome assembly

Contigs produced by ALLPATH-LG were optimized with the aid of contigs of PacBio assembly by using quickmerge with the parameters ‘-hco 5.0 -c 1.5 -l 100000 -ml 5000’. Optimized contigs were linked to scaffolds by leveraging Illumina mate-pair information by using SSPACE and gaps were filled by using PBjelly v2.

### Evaluation of oyster assembly

To appraise the genome quality, we first mapped Illumina reads to the oyster assembly by using Burrows-Wheeler Alignment (BWA) tool. Next, completeness of genomes was verified by mapping 248 highly conserved eukaryotic genes and 908 benchmarking universal single-copy orthologues in metazoa to the genomes by using CEGMA v2.5 [85] and BUSCO v3.0.2b [86], respectively.

### Hi-C sequencing and assembly

#### Sequencing

According to the Hi-C procedure [87], nuclear DNA from muscles of oyster individuals was cross-linked, then excised with a restriction enzyme, leaving pairs of distally located but physically intercalated DNA molecules attached to one another. The sticky ends of these digested fragments were biotinylated, which were then ligated to each other to form chimeric circles. Biotinylated circles, as chimeras of physically associated DNA molecules from the original cross-linking, were enriched, sheared and sequenced [88]. After adaptor removal and filtering out low-quality reads, Hi-C reads were aligned to our assembled genome to evaluate the ratios of mapped reads, distribution of insert fragments, sequencing coverage and number of valid interaction pairs. Uniquely mapped reads spanning two digested fragments that are distally located but physically associated DNA molecules are defined as valid interaction pairs.

#### Assembly

Scaffolds of PacBio+Illumina assembly were reduced to fragments with a length of 300 kb, which were then re-assembled by using the LACHESIS software [88] based on Hi-C data. Regions that failed to be restored to the original assembly or contained an average Hi-C data coverage of less than 0.5% were considered assembly errors, and were broken into smaller scaffolds. Consistency in assembly of Hi-C data based pseudo-chromosomes was assessed by comparisons with a genetic map for the *Crassostrea gigas* [89] by using software of ALLMAPS [90].

### Genome annotation

#### Repetitive sequence prediction

Repeat composition of the assemblies was estimated by building a repeat library employing the *de novo* prediction programs LTR-FINDER [91], MITE-Hunter [92], RepeatScout [93] and PILER-DF [94]. The database was classified by using PASTEClassifier [95] and then combined with the Repbase database [96] to create a final repeat library. Repeat sequences in oyster genome were identified and classified by using the RepeatMasker program [97]. The LTR family classification criterion was defined as that 5’-LTR sequences of the same family would share at least 80% identity over at least 80% of their lengths.

#### Protein-coding gene prediction

Protein-coding genes were predicted based on *de novo* and protein homology approaches. The algorithms Genscan [98], Augustus [99], GlimmerHMM [100], GeneID [101] and SNAP [102] were used for *de novo* gene prediction. Alignment of homologous peptides from *C. gigas*, *C. virginica*, *Lottia gigantea*, and *Danio rerio* to our assemblies was performed to identify homologous genes with the aid of GeMoMa [103]. Consensus gene models were generated by integrating the *de novo* predictions and protein alignments using EVidenceModeler (EVM) [104].

#### Functional annotation of protein-coding genes

Annotation of the predicted genes was performed by blasting their sequences against a number of nucleotide and protein sequence databases, including COG [105], KEGG [106], NCBI-NR and Swiss-Prot [107], with an *E*-value cutoff of 1e-5. Gene ontology (GO) for each gene were assigned by using Blast2GO [108] based on NCBI databases.

### Evolution of oysters

Protein sequences of *Haliotis discus hannai* [109], *Lottia gigantea* (GCF_000327385.1), *Aplysia californica* (GCF_000002075.1), *Biomphalaria glabrata* (GCF_000457365.1), *Crassostrea gigas* (GCF_000297895.1), *Crassostrea virginica* (GCF_002022765.2), *Pinctada fucata* (https://marinegenomics.oist.jp), *Chlamys farreri* (CfBase), *Bathymodiolus platifrons* (GCA_002080005.1), *Modiolus philippinarum* (GCA_002080025.1), *Octopus bimaculoides* (GCF_001194135.1), and *Homo sapiens* (GCF_000001405.26) were retrieved for analysis. Proteomes of the aforementioned twelve species and that of *C. hongkongensis*, comprising a total of 295,905 protein sequences, were clustered into 38,939 orthologue groups by using OrthoMCL v3.1 [110] based on an all-to-all BLASTP strategy with an *E*-value of 1e-5 and by using Markov Chain Clustering (MCL) algorithms with default inflation parameters (1.5). Based on clustering results, *C. hongkongensis*-specific gene families were determined and annotated. To infer phylogenetic relationships, we extracted 387 single-copy gene families from all thirteen species to perform multiple alignments of proteins for each family with MUSCLE v3.8.31 [111]. All of the alignments were combined into one supergene to construct a phylogenetic tree by using RAxML v8.2.12 [112] with 1000 rapid bootstrap analyses, followed by a search of the best-scoring ML tree in a single run. Finally, divergence times were estimated by using MCMCTree from the PAML package [113] in conjunction with a molecular clock model. Several reference-calibrated time points obtained from TimeTree database (http://timetree.org/) were used to date divergence times of interest. Expansion and contraction of OrthoMCL derived homologue clusters were determined by CAFÉ v2.1 [114] calculations on the basis of changes in gene family size with respect to phylogeny and species divergence time. In addition, we obtained domain-based expanded gene families of three *Crassostrea* species, according to previous works by *Albertin et al.* (2015) [115].

### Syntenic analysis

All-to-all BLASTP analyses of protein sequences were performed between *C. hongkongensis*, *C,gigas*, *C. virginica*, and *P. fucata* with an *E*-value threshold set at 1e-5. Syntenic regions within and between species were identified by using MCScan based on BLASTP results. A syntenic region was considered valid, if it contained a minimum of 10 collinear genes and a maximum of 25 gaps (genes) between two adjacent collinear genes.

### *Homeobox* gene analysis

Structures of *homeobox* genes in oyster were determined by using the GeMoMa v1.4.2 software [116] with default parameters based on available homeobox gene models. Predictions were handled by applying a GeMoMa annotation filter (GAF) with default parameters except for evidence percentage filter (e = 0.1). These were then manually verified to achieve a single high-confidence transcript prediction per locus. Exact annotations of each homeobox gene were completed with the aid of phylogenetic relationships.

### Transcriptomic analysis

Embryos at different developmental stages during oyster embryogenesis including zygote, 2-4 cells, blastula, morula, gastrula, trochophore, D-larva, veliger, pediveliger and spat were collected for RNA isolation. Similarly, RNA extraction was done with various tissues including hemocytes, muscles, gill, labial palp, hepatopancreas, gonads and mantles. To compare asymmetry-related mantle gene expression in the *C. hongkongensis* and *P. fucata*, their L/R mantles were collected. For both left and right mantles, unilateral tissues from five individuals were pooled as one sample, and each of the L/R mantle groups contained at least three replicates. Total RNA was isolated by using the Trizol reagent (Cat. no. 15596026, Invitrogen, CA), followed by treatment with RNase-free DNase I (Cat. no. M6101, Promega, WI), according to the manufacturers’ instructions. RNA quality was then checked by using an Agilent 2100 Bioanalyzer. Illumina RNA-Seq libraries were prepared and sequenced in a HiSeq 2500 system by a PE150 strategy following the manufacturer’s instructions (Illumina, CA). After trimming raw reads based on quality scores from the quality trimming program Btrim, clean reads were aligned to the oyster assembly genome by using TopHat v2.1.1 [117] and then assembled by using Cufflinks v2.1.1 [118]. Differential expression of genes in the various tissues was evaluated by using Cuffdiff [118].

### WGCNA and co-expression network analysis

Weighted correlation network analysis (WGCNA) [119] was applied to construct a weighted gene co-expression network of genes having a high correlation with cemented attachment. The top 10,000 differential genes exhibiting transcriptional changes in response to *L*-DOPA treatment were selected for WGCNA, wherein the modules showed high correlation with cemented attachment. We estimated the weight for each pair of genes forming intersections within these modules and analyzed differentially expressed genes relevant to cemented attachment by using DESeq2. Cytoscape [120] was used to delineate the co-expression network of significant gene pairs with weight >0.3.

### Byssal regeneration

Functional relationships between *antennapedia (Antp)* mRNA expression levels and phenotypic traits of byssal threads in adult pearl oysters (*Pinctada fucata*) were explored. Briefly, 50-100 pearl oysters (2 years old) were collected and maintained in aerated laboratory tanks. Byssal mass comprising the byssal stem and existing old threads of pearl oysters were excised. Then, individual pearl oysters were placed in beakers (one oyster per beaker) to allow identification of subsequent regrowth of nascent thread mass. Particular care was taken in removing old threads and attachment discs from the shells. Preliminary experiments indicate that removal of the threads did not affect subsequent thread formation. Byssal thread formation was estimated as the number of threads/oyster observed 24 h later.

Subsequently, the corresponding byssal gland of each pearl oyster was collected for RNA extraction by using TRIzol reagent, according to the manufacturer’s instructions. Purified RNA samples were diluted to 1 μg/μL and pooled to perform cDNA synthesis by utilizing PrimerScript first strand cDNA synthesis kit (Cat. no. 6110A, Takara Bio, Japan), following the manufacturer’s protocol. Real-time qPCR analysis was performed to determine *Antp* mRNA expression with gene-specific primers **(Table S16).**

### Pharmacological treatment

Chemical compounds were obtained from Sigma-Aldrich, unless otherwise specified. Working solutions were freshly prepared in deionized (DI) water approximately 1 h before *in vivo* experiments, which were conducted in large beakers to allow observation of oyster attachment and metamorphosis. Groups of oyster larvae at the pediveliger stage were placed in three beakers containing 50 mL sea water (at a density of 20 larvae/mL). There were three groups in total: an unstimulated control, an *L*-3,4-dihydroxyphenylalanine (*L*-DOPA) treatment and a norepinephrine (NE) treatment. Oyster larvae were challenged (6 h and 24 h) with different concentrations of NE (10^−4^, 10^−5^, 10^−6^ M) or *L*-DOPA (10^−5^, 10^−6^, 10^−7^ M). Previous studies have shown that this concentration range is sufficiently potent for inducing a larval response [121,122].

In addition, oyster larvae were collected following various treatment durations (6 h and 24 h) for RNA-seq and transcriptomic analysis to determine any temporally driven differences between the *L*-DOPA treatment group (10^−5^ M) and unstimulated control. By a similar design, oyster larvae were exposed to NE (10^−5^ M) for 6 h and 24 h, and their transcriptomic profiles were examined in relation to oyster metamorphosis.

## Data availability

The *C. hongkongensis* genome studied in this Hong Kong oyster genome project has been deposited at the NCBI under the BioProject number PRJNA592306 at https://www.ncbi.nlm.nih.gov/bioproject/PRJNA592306. Hi-C data have been deposited as SRR10583824 at https://www.ncbi.nlm.nih.gov/sra/SRR10583824. RNA-seq data of various transcriptomes have been deposited as PRJNA588628 at https://www.ncbi.nlm.nih.gov/bioproject/PRJNA588628.

## CRediT author statement

**Yang Zhang**: Conceptualization, Methodology, Validation, Investigation, Data Curation, Writing - Original Draft, Writing - Review & Editing, Visualization, Supervision, Project administration, Funding acquisition. **Fan Mao**: Methodology, Validation, Investigation, Data Curation, Writing - Original Draft, Writing - Review & Editing, Visualization, Funding acquisition. **Shu Xiao**: Methodology, Validation, Resources, Funding acquisition. **Haiyan Yu**: Methodology, Formal analysis, Investigation, Data Curation. **Zhiming Xiang**: Methodology, Validation, Data Curation, Funding acquisition. **Fei Xu**: Formal analysis, Validation, Data Curation. **Jun Li**: Validation, Resources. **Lili Wang**: Formal analysis. **Yuanyan Xiong**: Formal analysis. **Mengqiu Chen**: Formal analysis. **Yongbo Bao**: Formal analysis. **Yuewen Deng**: Validation. **Quan Huo**: Validation. **Lvping Zhang**: Validation. **Wenguang Liu**: Validation. **Xuming Li**: Formal analysis. **Haitao Ma**: Formal analysis. **Yuehuan Zhang**: Resources. **Xiyu Mu**: Formal analysis. **Min Liu**: Formal analysis. **Hongkun Zheng**: Conceptualization, Formal analysis, Data Curation, Project administration. **Nai-Kei Wong**: Writing - Review & Editing, Visualization. **Ziniu Yu**: Conceptualization, Writing - Review & Editing, Visualization, Supervision, Project administration, Funding acquisition.

## Competing interest

We declare that none of the authors have competing financial or non-financial interests.

## Acknowledgments

We are deeply grateful to our lab members and collaborators, who have provided us with able assistance or valuable advice at all stages of this study. We acknowledge grant support from Key Special Project for Introduced Talents Team of Southern Marine Science and Engineering Guangdong Laboratory (Guangzhou) (GML2019ZD0407), Key Deployment Project of Centre for Ocean Mega-Research of Science, Chinese Academy of Science (COMS2019Q11), the National Science Foundation of China (No. 32073002, 31902404), the China Agricultural Research System (No. CARS-49), the Science and Technology Program of Guangzhou, China (No.201804020073), Natural Science Foundation of Guangdong Province (2020A1515011533), the Program of the Pearl River Young Talents of Science and Technology in Guangzhou of China (201806010003), Institution of South China Sea Ecology and Environmental Engineering, Chinese Academy of Sciences (ISEE2018PY01, ISEE2018PY03, ISEE2018ZD01), and Science and Technology Planning Project of Guangdong Province, China (2017B030314052, 201707010177).

## Notes

### Competing Interest Statement

The authors have declared no competing interest.

