## supplemental Figure 1-15 for "Comparative genomics reveals evolutionary drivers of sessile life and left-right shell asymmetry in bivalves": Supplementary Figures for preprint.docx

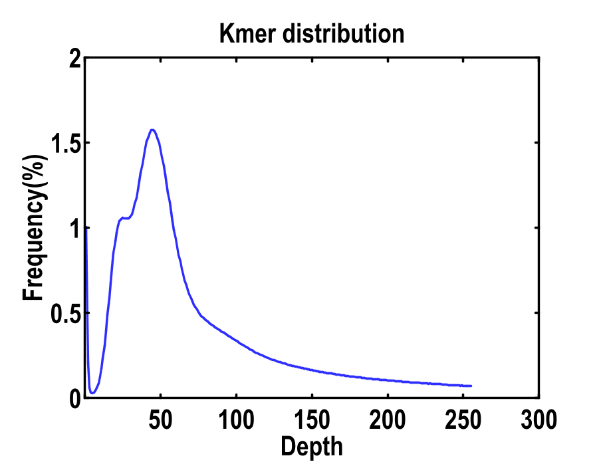

**Figure S1: Seventeen k-mer depth distribution of *Crassostrea hongkongensis* genome sequences.** Depth distribution of k-mer showed a major peak at 45×, and genome size was estimated to be ~593 Mb according to the formula: Genome size= k-mer number/peak depth.

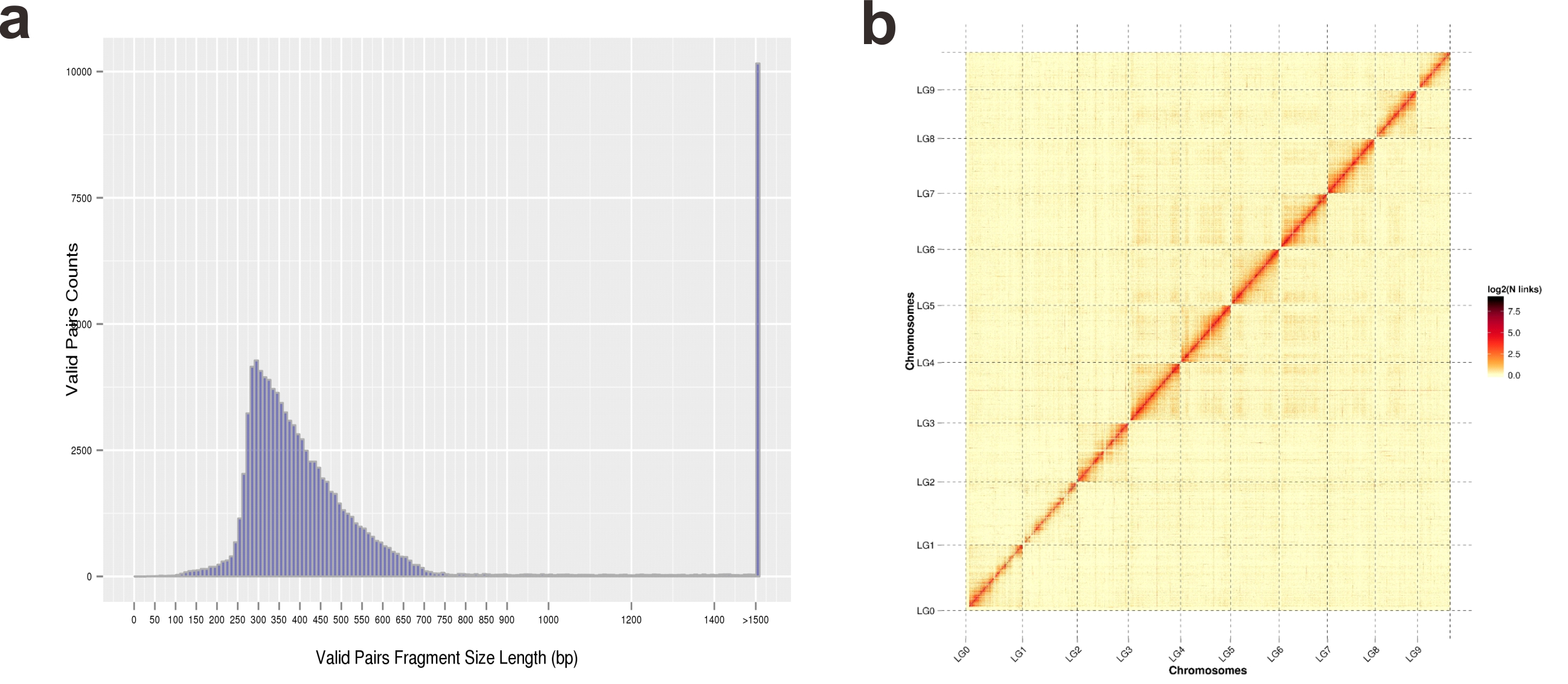

**Figure S2a: Count distribution of valid-pairs fragment size.** A total of 36.82 Gb clean Hi-C reads were aligned to the *C. hongkongensis* genome assembled genome. Valid interaction pairs were defined as unique mapped reads spanning two digested fragments, which were distally located but physically associated DNA molecules. **Figure S2b: Chromosomal *C. hongkongensis* genome was assembled with the aid of Hi-C data.**

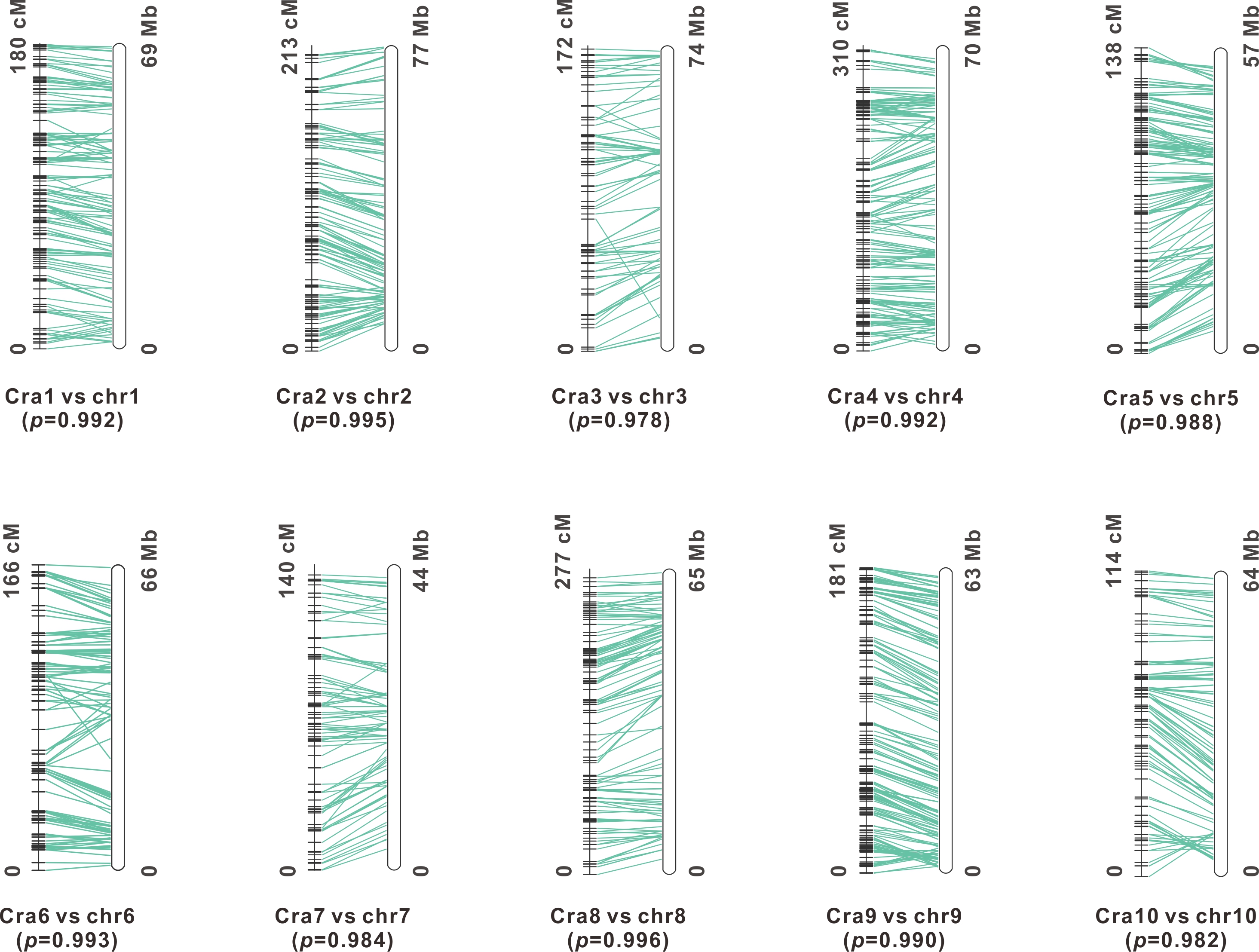

**Figure S3: Consistency between Hi-C data based pseudo-chromosomes and a genetic map for the relative species, *Crassostrea gigas***. Chr1-10 and Cra1-10 indicate the Hi-C data based pseudo-chromosomes of *C.hongkongensis* and the genetic map for *Crassostrea gigas*, respectively*. p* values refer to their consistency, which was assessed by using the software ALLMAPS.

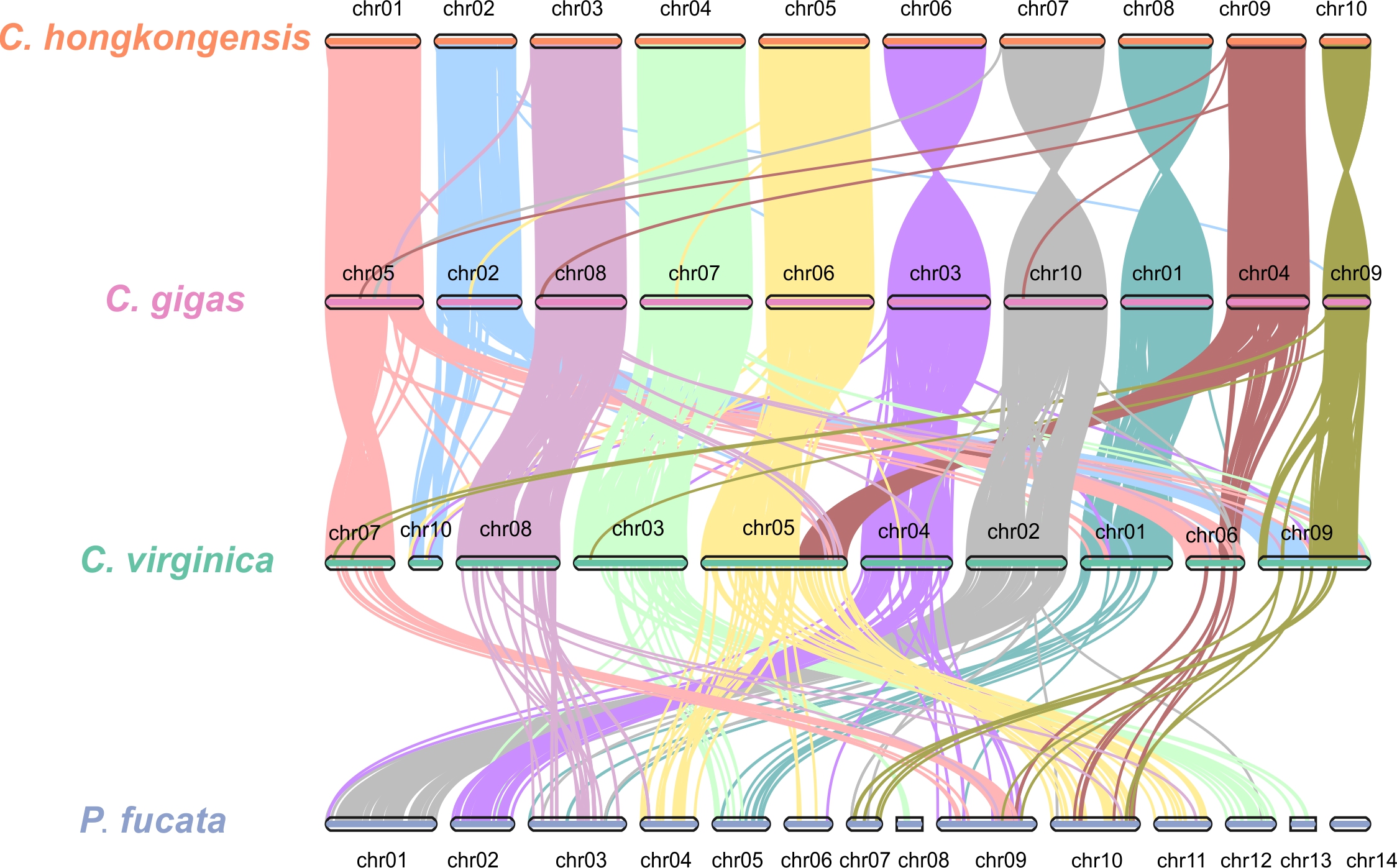

**Figure S4: Genomic synteny between the genomes of three *Ostreoida* oysters and *Pinctada fucata*.** All-to-all BLASTP analyses of protein sequences were set at threshold of 1e-5. Syntenic regions within and between species were identified by using MCScan based on BLASTP results. A syntenic region was defined as a minimum of 10 collinear genes and a maximum of 25 gaps (genes) between two adjacent collinear genes.

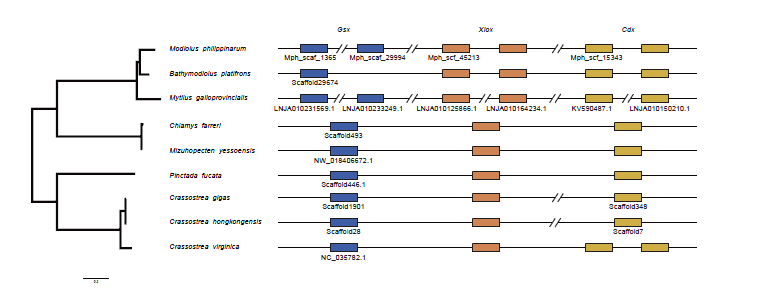

**Figure S5: Evolution and organization of the paraHox gene cluster in bivalves.** A paraHox cluster composed of three genes, *Gsx, Xlox and Cdx*. Schematic representation of paraHox genomic evolution and organization in several bivalves: *Modiolus philippinarum*, *Bathymodiolus platifrons*, *Mytilus galloprorinclas*, *Mytilus galloprorinclas*, *Mizuhopecten yessoensis*, *Pinctada fucata*, *Crassostrea gigas*, *Crassostrea hongkongensis*, and *Crassostrea virginica*. Boxes indicate paraHox genes in the genome. A continuous line indicates an intact cluster. Double-parallel lines indicate long genomic distances.

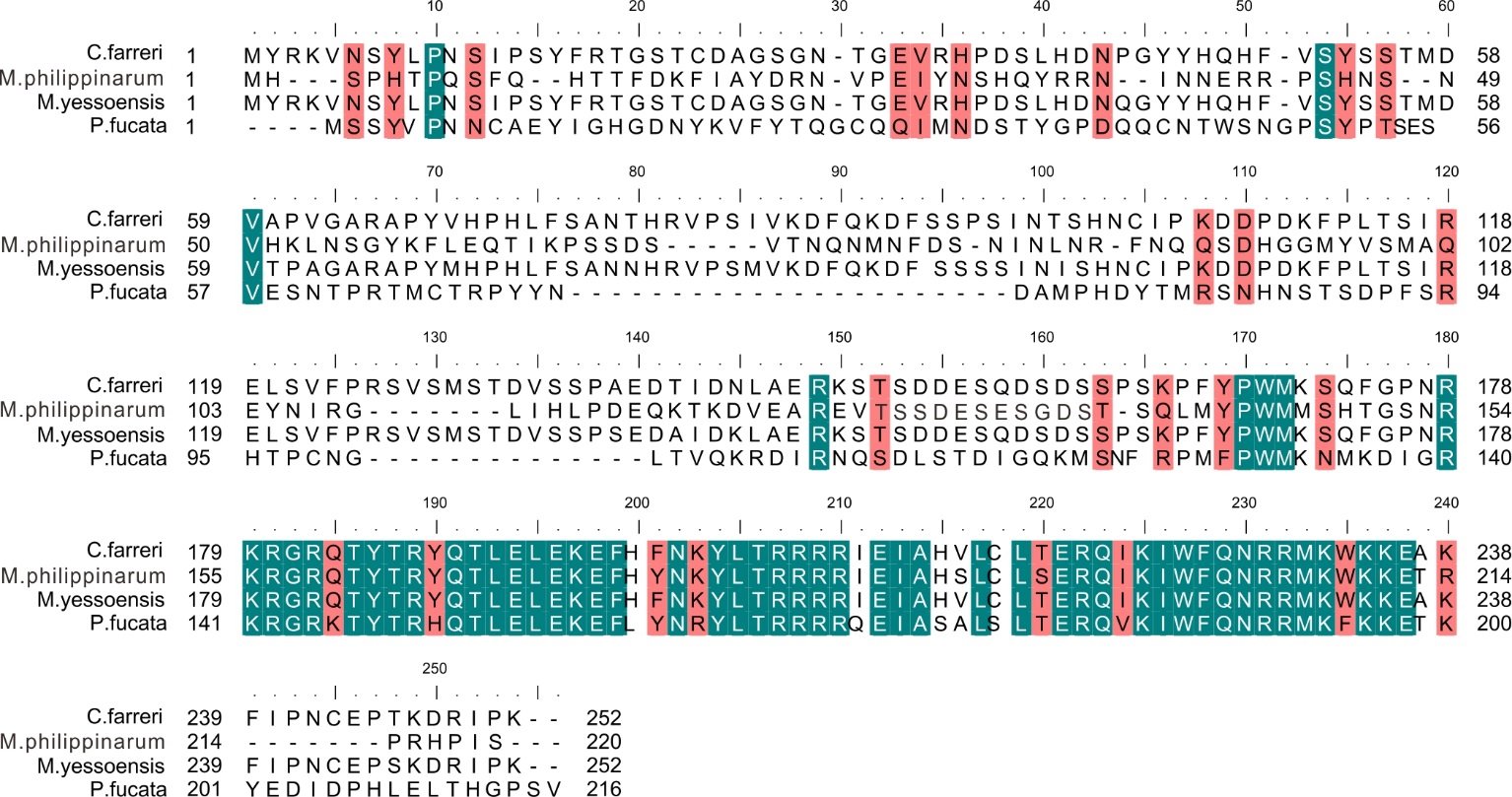

**Figure S6: Sequence al****ignment of *antennapedia* (*Antp*) amino acid sequences.** Amino acid sequences of four byssally attached bivalve species (*Chlamys farreri*, *Modiolus philippinarum*, *Mizuhopecten yessoensis*, and *Pinctada fucata*) were obtained and aligned by using the Bioedit software. A conserved region with high identity, the Hox domain, can be observed in these four byssally attached bivalve species, which is known to be involved in transcriptional regulation of key developmental processes. In the figure, red color marks the sites with high sequence similarity, while green marks the sites with the same sequence.

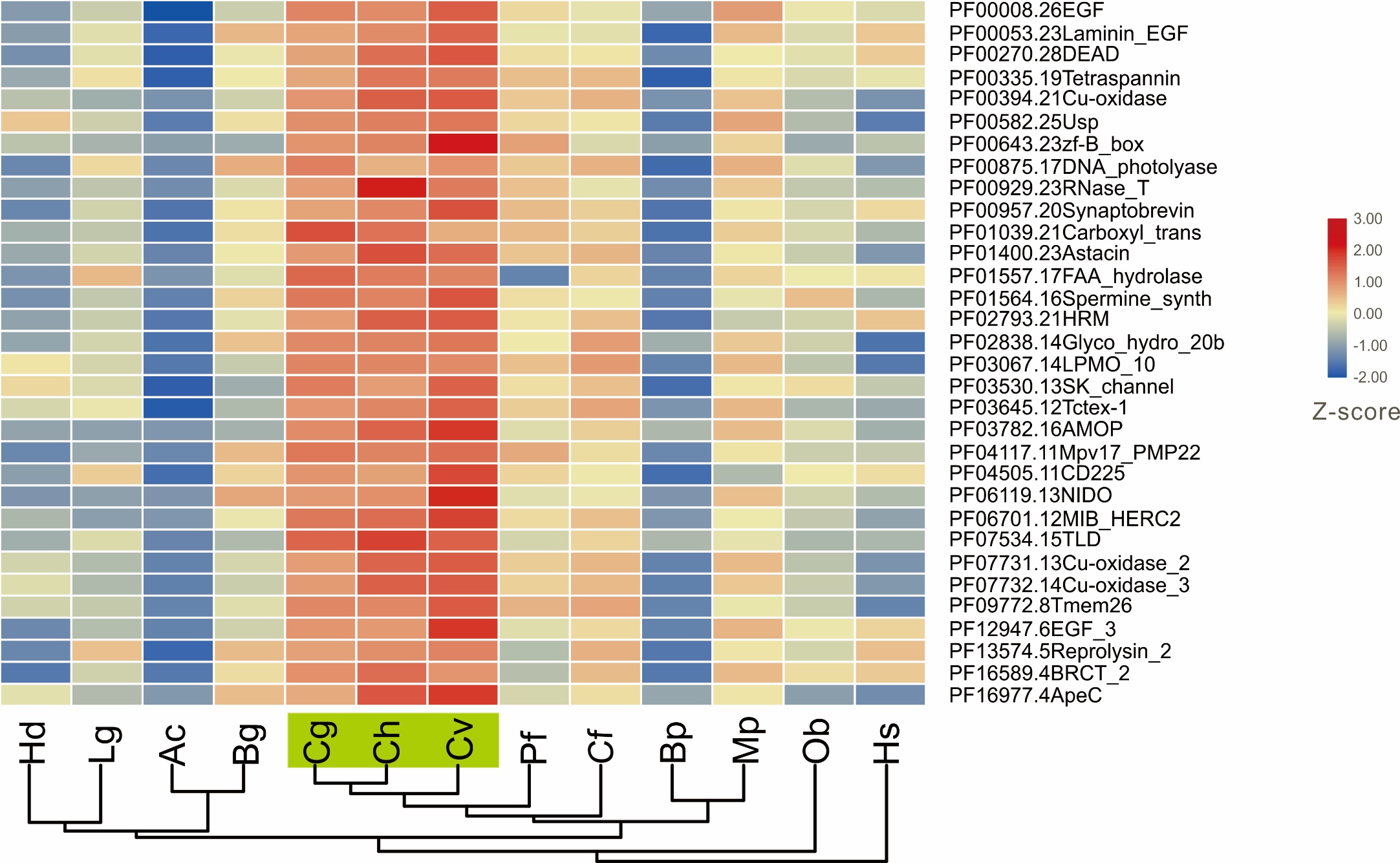

**Figure S7: Heatmap presents 32 of common expanded gene families.** A heatmap was generated using TBtool software(1) according to the gene number of each gene family. Evidently, these gene families are greatly expanded in three *Ostreoida* oysters under study, namely, *C. gigas*, *C. hongkongensis*, and *C. virginica*. *Haliotis discus hannai* (Hd), *Lottia gigantea* (Lg), *Aplysia californica* (Ac), *Biomphalaria glabrata* (Bg), *Crassostrea gigas* (Cg), *Crassostrea hongkongensis* (Ch), *Crassostrea virginica* (Cv), *Pinctada fucata* (Pf), *Chlamys farreri* (Cf), *Bathymodiolus platifrons* (Bp), *Modiolus philippinarum* (Mp), *Octopus bimaculoides* (Ob), and *Homo sapiens* (Hs).

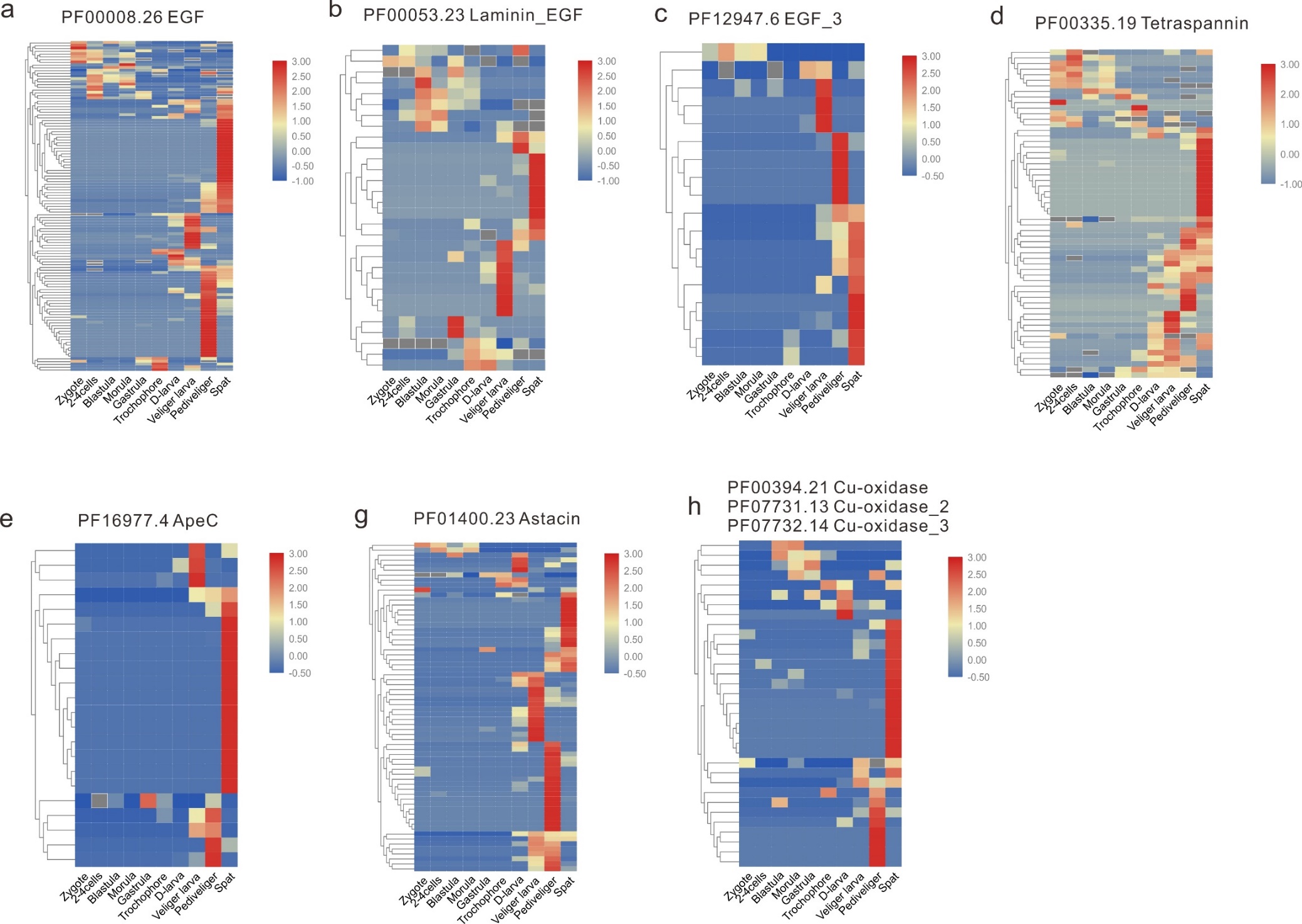

**Figure S8: Heatmaps of high-correlation gene families in oyster larval attachment.** Gene families of high correlation with larval attachment across developmental stages were analyzed, including *PF00008.26 EGF* (**a**)*, PF00053.23 Laminin_EGF* (**b**)*, PF12947.6 EGF_3* (**c**)*, PF00335.19 Tetraspannin* (**d**)*, PF16977.4 ApeC* (**e**)*, PF02793.21 HRM* (**f**)*, PF01400.23 Astacin* (**g**)*,* and *PF00394.21*, *PF07731.13*, and *PF07732.4 Cu-oxidase* (**h**). Changes in gene expression level during different developmental stages are schematically depicted. Heatmaps were generated using TBtool software(1). Developmental stages are abbreviated as: zygote, 2-4 cells, blastula, morula, gastrula, trochophore, D-larve, veliger larva, pediveliger, and spat.

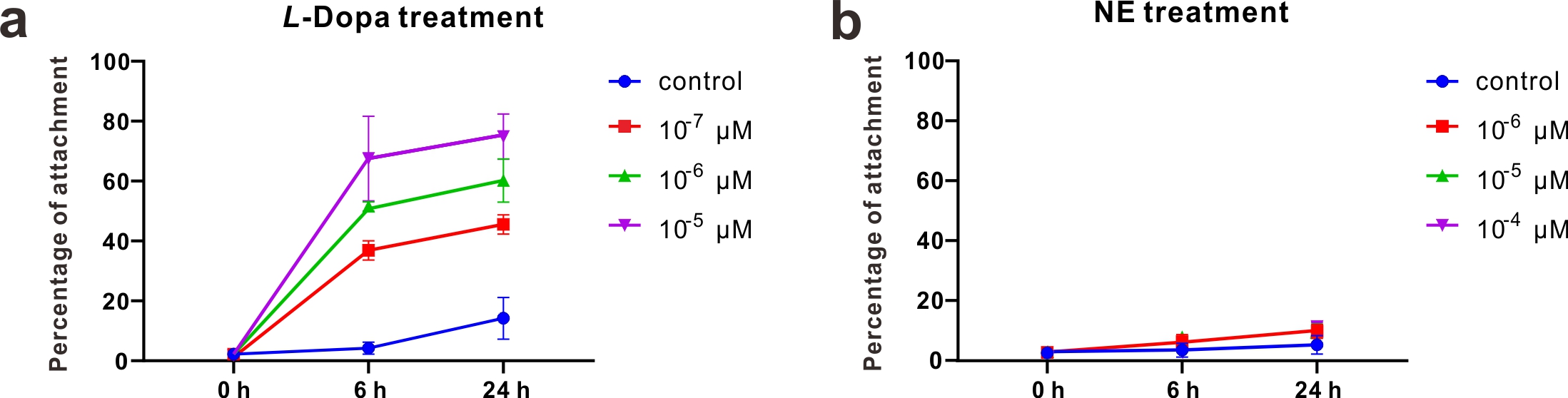

**Figure S9: Effects of *L*-DOPA and NE treatments on oyster larval attachment.** *L*-3,4-dihydroxyphenylalanine (*L*-DOPA) induced normal attachment and metamorphosis, whereas noradrenaline (NE) induced metamorphosis without attachment. Low-concentration (10^-6^ M) *L*-DOPA treatment caused an increase in adhesion rate, while medium- (5$\times$10^-6^ M) and high-concentration (10^-5^ M) treatments caused an increase in adhesion rate within a short time (3 h). NE induced morphological changes in larvae, and treatment at 3 h and 6 h significantly induced metamorphosis. There was no significant induction effects after 9 h treatment. We also determined the survival rate of oyster larvae after 30 h of drug treatment. Results show that all experimental concentrations used in drug treatment did not affect larval survival rate compared to that of the control group.

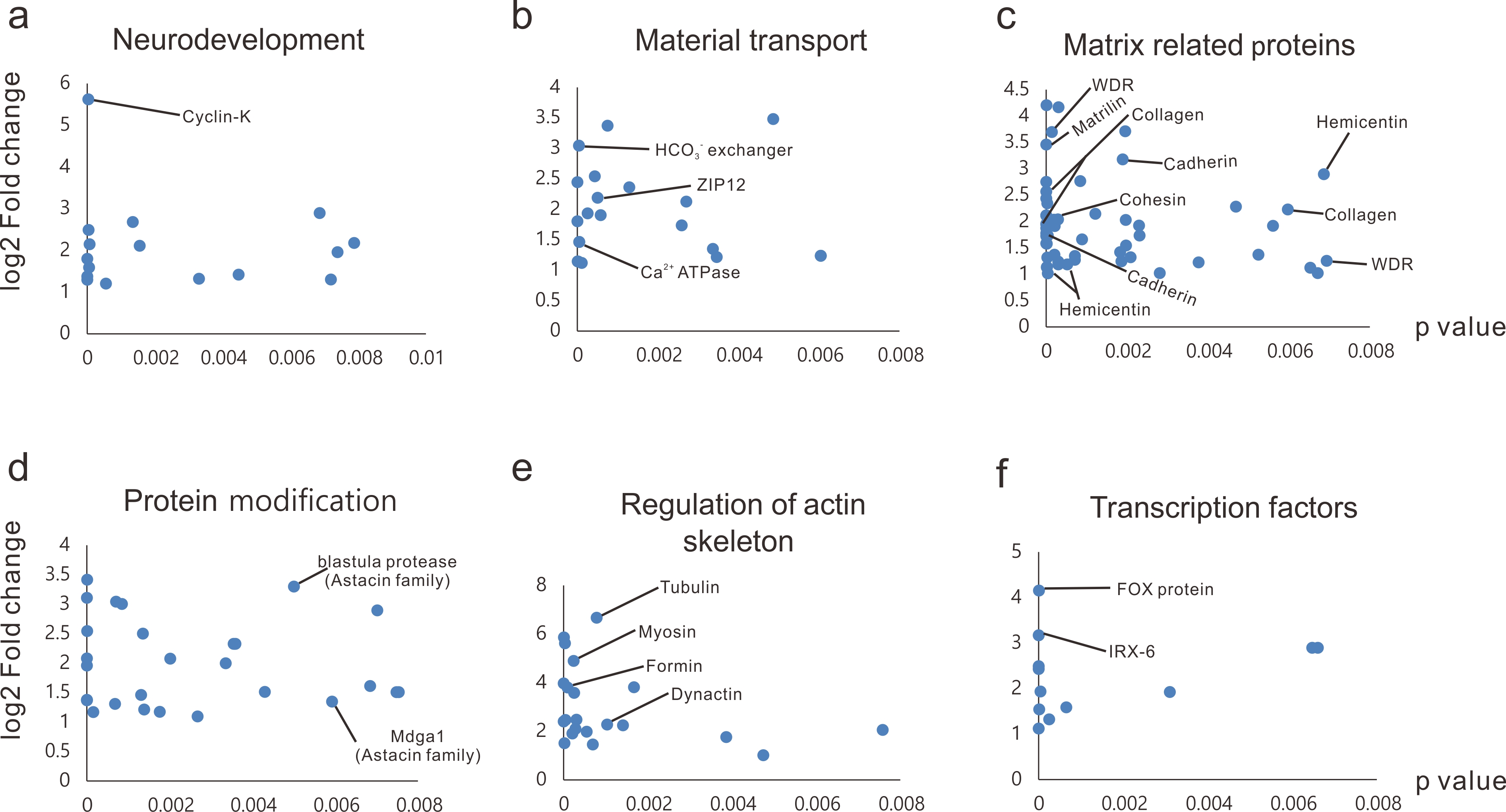

**Figure S10: Transcriptomic analysis post-treatment.** Transcriptome sequencing was conducted to assess specific expression of 1,424 genes after *L*-DOPA treatment, involved in neurodevelopment, material transport, matrix related proteins, protein modification, regulation of actin skeleton, and transcription factors. Some important genes are marked in the figure. The *x*-axis represents *p* values and the *y*-axis represents fold changes.

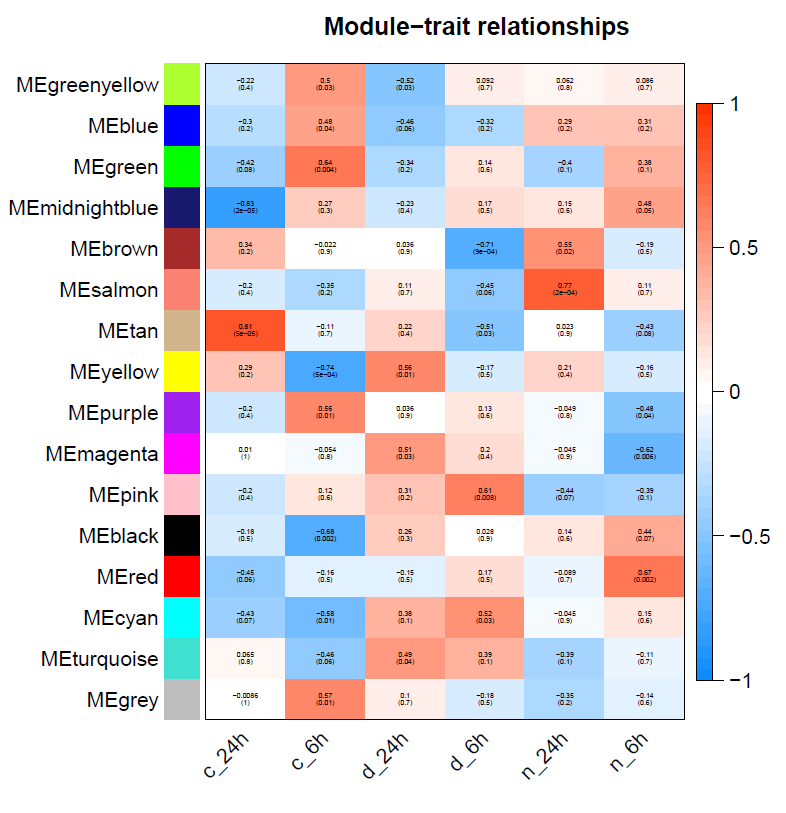

**Figure S11:** **WGCNA analysis on module-trait relationships.** The top 10,000 differentially expressed genes exhibiting transcriptional changes in response to *L*-DOPA treatment were selected for WGCNA, wherein MEpink modules showed a high correlation with *L*-DOPA induced attachment.

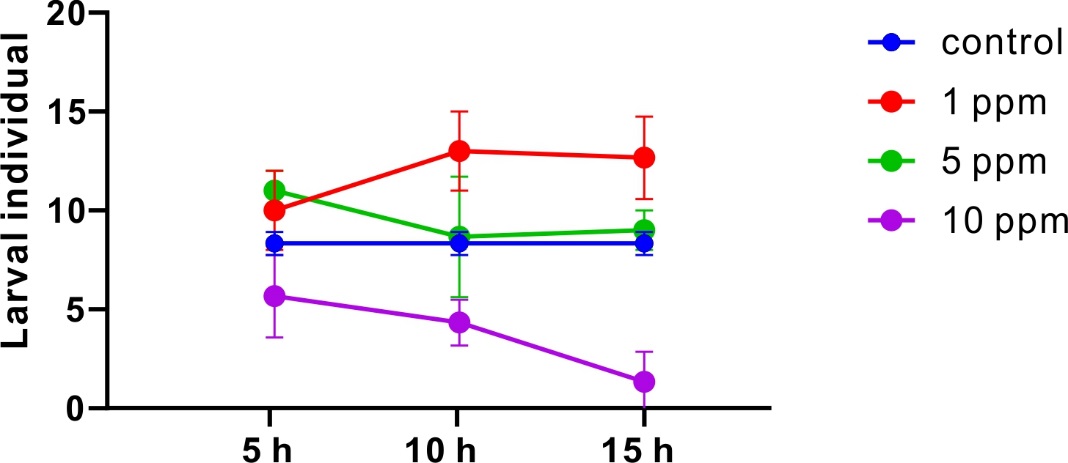

**Figure S12: Low-dose zinc induces attachment.** Oyster larvae were treated with different concentrations of ZnCl_2_ (1 ppm, 5 ppm, and 10 ppm) for 5 h, 10 h, and 15 h. The control groups were treated with equal volumes of sea water. Each group contained three replicates and independent experiments were performed three times. Error bars represent means $\pm$ SEM (*n* $=$3).

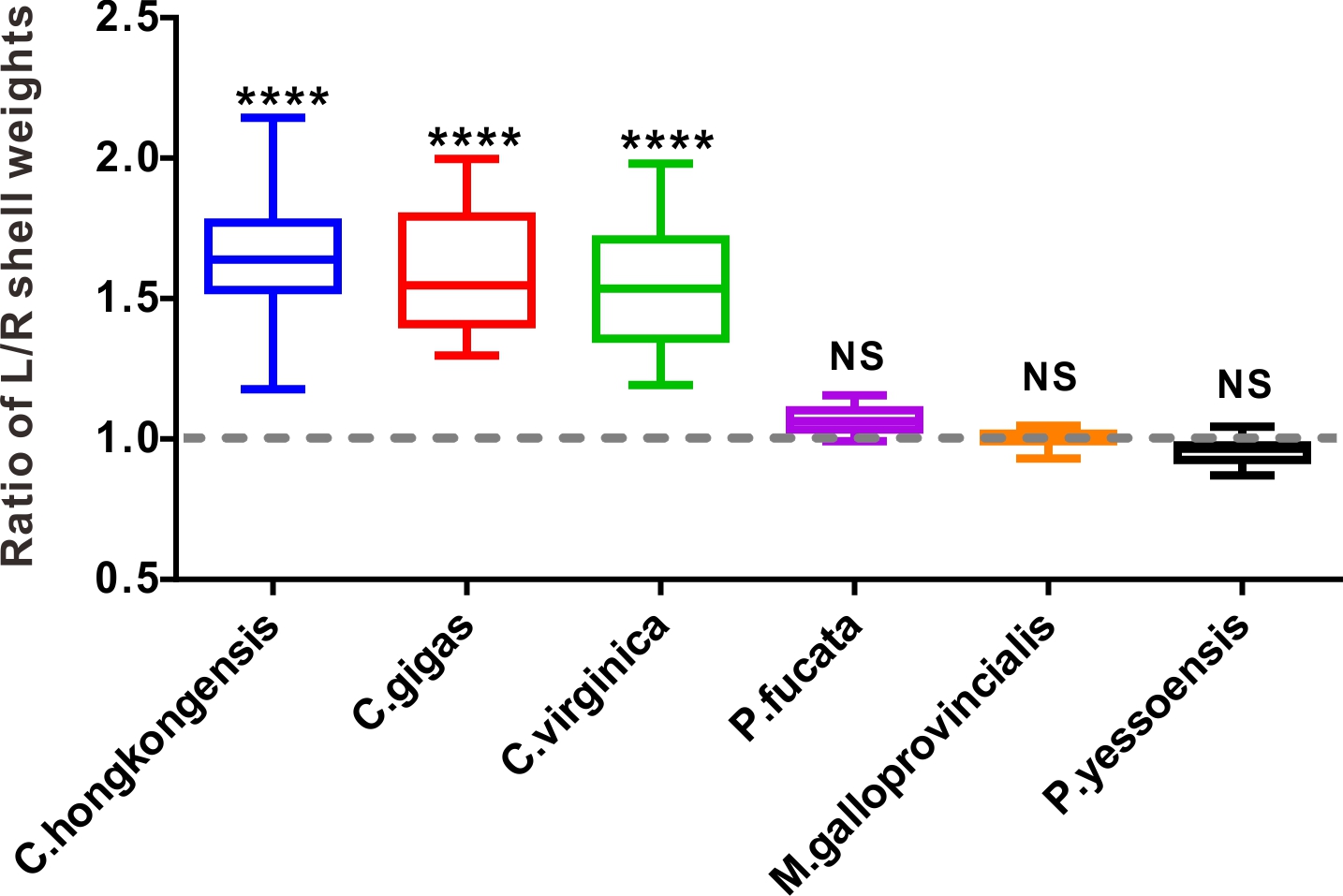

**Figure S13: Ratio of L/R shell weights in different bivalve species.** Weights of the left and right shells of several representative bivalves were measured and their statistical results are as shown in the figure. Forty individuals per species were used for the assessment. The left and right shell weights of six bivalves are represented in a box plot. Dotted line indicates a weight ratio of 1. ****, *p* <0.0001.

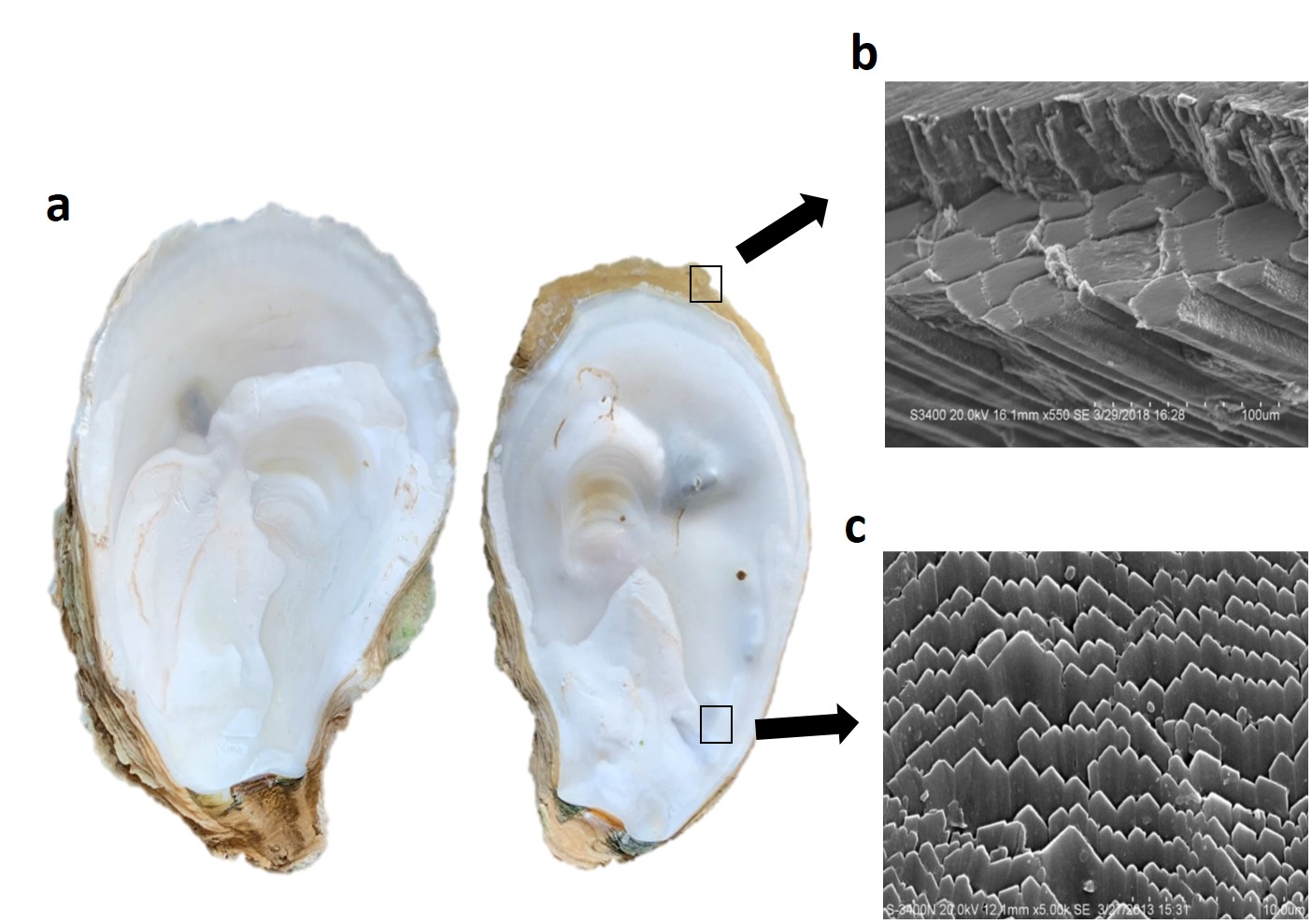

**Figure S14: Scanning electron micrographs of the left and right shells of the Hong Kong oyster.** Oyster shells consist of a prismatic layer and a foliate layer spanning from outside to inside and are chemically composed of CaCO_3_. **a,** The left and right shell of the Hong Kong oyster. **b,** The top side of right oyster is the prismatic layer. c, The inner side of right shell is the foliate layer. SEM magnified the object 550 times (b) and 5,000 times (c).

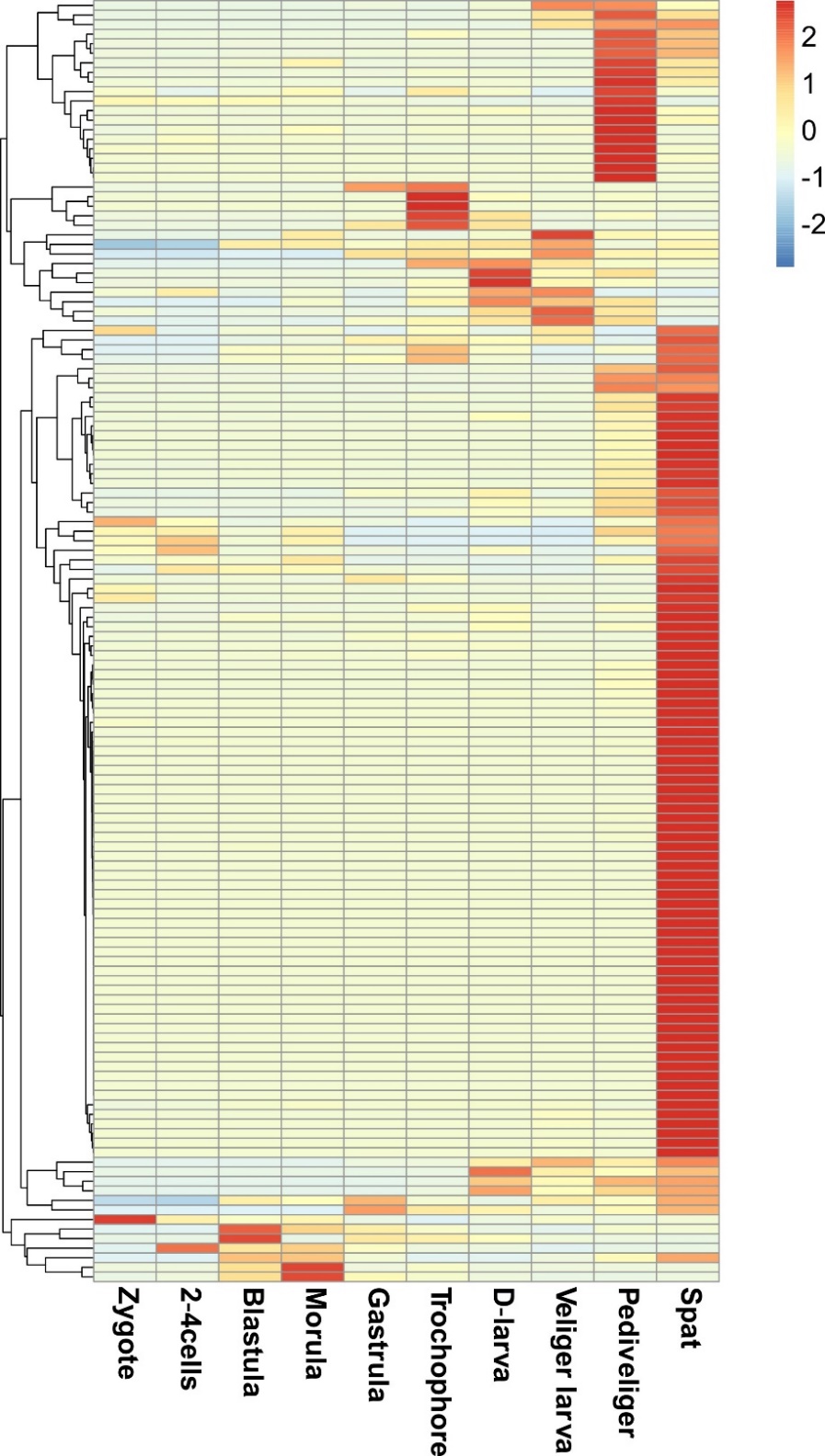

**Figure S15: Shell asymmetry-related gene expression profiles seen during the developmental stages.** To study the molecular mechanisms underlying asymmetric growth of oyster shells in the Hong Kong oyster, expression patterns of 134 differentially expressed genes were analyzed during developmental stages. Developmental stages are abbreviated as: zygote, 2-4 cells, blastula, morula, gastrula, trochophore, D-larve, veliger larva, pediveliger, and spat.
